# Drivers of mangrove vulnerability and resilience to tropical cyclones in the North Atlantic Basin

**DOI:** 10.1101/2022.11.22.517275

**Authors:** Cibele Hummel do Amaral, Benjamin Poulter, David Lagomasino, Temilola Fatoyinbo, Paul Taillie, Gil Lizcano, Steven Canty, Jorge Alfredo Herrera Silveira, Claudia Teutli-Hernández, Miguel Cifuentes, Sean Patrick Charles, Claudia Shantal Moreno, Juan David González-Trujillo, Rosa Maria Roman-Cuesta

## Abstract

The North Atlantic Basin (NAB) is seeing a significant increase in the frequency and intensity of tropical cyclones since the 1980s, with record-breaking seasons such as 2017 and 2020. However, little is known about how coastal ecosystems, particularly mangroves in the Gulf of Mexico and the Caribbean, are responding to these new “climate normals” at regional and subregional scales. Wind speed, rainfall, pre-cyclone forest structure, and hydro-geomorphology are known to influence mangrove damage and recovery following cyclones in the NAB. However, these studies have focused on site-specific responses and individual cyclonic events. Here, we analyze 25 years (1996-2020) of mangrove vulnerability (damage after a cyclone) and short-term resilience (recovery after damage) for the entire NAB and its subregions, using multi-annual, remote sensing-derived databases. We applied machine learning to characterize the influence of 22 potential drivers that include previously researched variables and new ones such as human development and long-term climate trends. The characteristics of the cyclones mainly drive vulnerability at the regional level, while resilience is largely driven by site-specific conditions. These include long-term climate conditions, such as air temperature and drought trends, pre-cyclone habitat conditions, such as canopy cover and height and soil organic carbon stock, and human interventions on the land. Rates and drivers of mangrove vulnerability and resilience vary across subregions in the NAB, and hotspots for restoration and conservation actions are highlighted within subregions. The impacts of increasing cyclone activity need to be framed in the context of climate change compound effects and heavy human influences in the region. There is an urgent need to value the restoration and conservation of mangroves as fundamental Nature-based Solutions against cyclone impacts in the NAB.

## 1. Introduction

Worldwide, mangrove forests buffer coastal areas and their economies from wind and storm surges (Hochard et al., 2019; Menéndez et al., 2020; Zhu et al., 2020), while also storing carbon at some of the highest densities of any ecosystem (Donato et al., 2011; Richards et al., 2020). Globally, their coastal protective benefits are estimated to avert US$ 60 billion in flooding damages from tropical cyclones and shield 14 million people from their devastating impacts (Menéndez et al., 2020). Flooding benefits vary among regions and countries due to different exposures, mangrove extents, and cyclone characteristics, but NAB countries top the list in averted land flooding and reduced damages to people and property due to mangrove presence (Menéndez et al., 2020). While efforts are growing to internalize the Nature-based Solutions (NbS) provided by mangroves into innovative financial mechanisms that may help protect them (Earth Security, 2020), their conservation seems to remain untapped in regional policies on coastal planning, and disaster risk reduction in the NAB. Simultaneously, the risk of tropical cyclones is predicted to increase in the region (Bacmeister et al., 2018; Emanuel, 2021; Knutson et al., 2021) where higher cyclone frequencies and intensities are already leading to enhanced coastal damages (Wang & Tuomi, 2021) reaching more exposed and increasingly more vulnerable societies (Hsiang & Jina, 2014; ECLA, 2018; Ötker & Srinivasan, 2018; Ramenzoni et al., 2020).

While mangroves are known to be resilient to the long-term impacts of tropical cyclones (Lugo, 1980; Jimenez et al., 1985; Roth, 1992; Kraus & Osland, 2020) and can even benefit from a storm’s nutrient influx (Castañeda-Moya et al., 2020), there is increasing post-cyclone mortality that may signal the presence of tipping points (Taillie et al., 2020; Lagomasino et al., 2021). This was the case of the 2017 Mega-Hurricane season, when abnormal mangrove mortality was observed a year after the hurricane season, resulting in 30-times more mangrove mortality than any year during the period 2009-2018 (Taillie et al., 2020). Reasons behind this unexpected mortality event remain unclear. Although, compounded factors such as the preceding, severe, and long-lasting El Nino event in the region (2015-2016), have been suggested as a yet under-recognized but deadly “drought-hurricane duo”, as it has been reported with compounded extreme events in other tropical forests (Brando et al., 2014; Allen et al., 2021; Berenguer et al., 2021). Moreover, the nature of tropical cyclones is varying in the NAB with more erratic and stalling storms that bring significantly higher amounts of rainfall (Hall & Kossin, 2019), whose impounding effects (drowning) are less understood in leading to mangrove mortality than the impacts of wind uprooting and sediment displacement (Roth, 1992; Imbert, 2018; Kraus & Osland, 2020; Lagomasino et al., 2021).

Since more frequent damage to mangroves from tropical cyclones compromise the buffering capacity of mangroves to future cyclonic events (Danielson et al., 2017), it is imperative to identify where NAB regional hotspots of cyclone-driven mangrove damage and loss of resilience exist, so that national managers and regional policymakers can act on their long-term restoration and conservation. It is also critical to identify the drivers behind widespread mangrove damage and loss of resilience after tropical cyclones, in order to define the best management for each case while enhancing mangrove functionality and persistence (Gijsman et al., 2021). In this line, previous research has shown that mangroves in the NAB are more vulnerable to cyclones if they are structurally taller than surrounding trees, wind speeds are ≥100 km h^-1^, storms bring high flooding, cyclone recurrence is relatively low (only affected by a few cyclones before), and the composition is dominated by red mangrove (*Rhizophora mangle* L.) (Smith, 2009; Imbert, 2018; Sippo et al., 2018; Rivera-Monroy et al., 2020; Taillie et al., 2020; Lagomasino et al., 2021; Peereman et al., 2022) On the other hand, mangroves exhibit lower short-term resilience after being damaged if they are located in poorly drained basins that might promote impounding and salinity increases, in areas with lower fertility, or where their composition is dominated by black or white mangrove (*Avicennia germinans* (L.) L., *Laguncularia racemosa* (L.) C.F.Gaertn.) (Smith, 2009; Harris et al., 2010; Vogt et al., 2012; Imbert, 2018; Rivera-Monroy et al., 2020, Lagomasino et al., 2021).

While these studies offer good insights on mangrove vulnerability and resilience to tropical cyclones in the NAB, they 1) have only captured the responses on specific sites and/or one-year post-disturbance responses and/or 2) have mainly included environmental variables and short-term climate responses without assessing long-term climate trends and means and human drivers such as the influence of urbanization, croplands, and road networks, which are also known to act as drivers of mangrove degradation and loss (Feller et al., 2015; Hayashi et al., 2019; Branoff, 2020; Goldberg et al., 2020; Villate Daza et al., 2020). Nor have these previous studies captured the difference in vulnerability and resilience of mangroves to cyclones at subregional levels in the NAB. We address these research gaps and policy-relevant questions using freely-available spatial data. Here, we specifically pose three major questions: 1) To what degree have mangroves in the NAB been affected by cyclones (from tropical storms to hurricanes) in the period 1996-2020? 2) What are the most influential human and environmental drivers behind mangrove damage (vulnerability) and recovery (resilience) to cyclones in the NAB? 3) How do drivers vary between different subregions? Solving those questions is of interest for risk assessments, adaptation policies, and ecosystem restoration and conservation in the NAB region.

## 2. Material and Methods

### 2.1. Study site

The study was carried out in the NAB, specifically in the Caribbean and Gulf of Mexico where mangroves occur. The study site covers an area of 7,741,775.5 km^2^ from South to North America (i.e., 7-30°N and 60-98°W), hosts 35 countries, and can be subdivided into nine coastal ecoregions, according to Spalding et al. (2007) (Table 1 and **Fig. 1**). This bioregionalization was created to support national and international conservation policy agendas and are based on global biogeographic patterns (Spalding et al., 2007). In 1996, the first year of our study, the NAB had 2,025,295 ha of mangroves (Bunting et al., 2018), with the Southern Gulf of Mexico and the Greater Antilles hosting the largest extents of mangroves and the Northern Gulf of Mexico the smallest (Table 1).

**Table 1.** Study site description by subregion (Spalding et al., 2007): countries, mangrove area (ha) for the year 1996 (Bunting et al., 2018), cumulative maximum wind speed, and hurricane season rainfall from 1980 to 2020 (Hersbach et al., 2018), mean annual maximum temperature, mean annual rainfall, and trend of the Self-calibrating Palmer Drought Severity Index (scPDSI) from 1982 to 2016 (CRU et al., 2017), and the number of settlements (CIESEN, 2011) and length of roads (CIESEN, 2013) within 5 km from the coast.

| Subregion | Countries | Mangrove area (ha) | Cyclonic Activity |  | Climate |  |  | Human Infrastructure |  |
| --- | --- | --- | --- | --- | --- | --- | --- | --- | --- |
|  |  |  | Cumulative Max. Wind Speed (km/h) | Cumulative hurricane season rainfall (mm) | Annual max. temperature (°C) | Annual rainfall (mm) | Drought trend | Settlements (n) | Roads (km) |
| **Bahamian** | The Bahamas, Turks and Caicos Islands (UK) | 167,475 | 2875.4 ± 200.7 | 29134.6 ± 4042.4 | 30.0 ± 0.9 | 105.2 ± 6.7 | -0.0523 ± 0.0302 | 38 | 1,782.2 |
| **Eastern Caribbean** | American Virgin Islands (US), Anguilla (UK), Antigua and Barbuda, Barbados, British Virgin Islands (UK), Dominica, Grenada, Guadeloupe (FR), Martinique (FR), Montserrat (UK), St. Kitts and Nevis, St. Lucia, St. Vincent, and the Grenadines | 87,945 | 2480.9 ± 199.5 | 29830.00 ± 4315.3 | 28.6 ± 1.3 | 197.5 ± 45.0 | 0.0123 ± 0.0402 | 201 | 3,466 |
| **Floridian** | United States | 243,409 | 2670.9 ± 295.4 | 30913.8 ± 3626.2 | 28.5 ± 0.4 | 113.0 ± 7.8 | -0.0003 ± 0.0401 | 313 | 4,308 |
| **Greater Antilles** | Cayman Islands (UK), Cuba, Dominican Republic, Haiti, Jamaica, Puerto Rico (US), United States | 489,790 | 2467.2 ± 269.9 | 29889.6 ± 6019.1 | 29.7 ± 1.2 | 127.9 ± 30.5 | 0.0211 ± 0.0413 | 210 | 8,819 |
| **Northern Gulf of Mexico** | Mexico, United States | 12,098 | 2671.0 ± 347.3 | 27461.3 ± 5283.6 | 27.2 ± 1.6 | 88.1 ± 32.5 | -0.0121 ± 0.0258 | 358 | 7,148 |
| **Southern Caribbean** | Aruba (NL), Bonaire (NL), Colombia, Curacao (NL), Trinidad and Tobago, Venezuela | 52,552 | 1928.2 ± 500.7 | 39036.0 ± 24285.7 | 31.1 ± 2.4 | 104.3 ± 32.1 | -0.0116 ± 0.0319 | 45 | 3,148 |
| **Southern Gulf of Mexico** | Guatemala, Mexico | 559,204 | 2486.1 ± 524.7 | 33822.1 ± 14194.1 | 29.5 ± 2.9 | 117.2 ± 50.0 | 0.0106 ± 0.0445 | 44 | 6,800 |
| **Southwestern Caribbean** | Colombia, Costa Rica, Honduras, Nicaragua, Panama, Venezuela | 175,322 | 2174.7 ± 523.8 | 60275.8 ± 23567.5 | 30.3 ± 2.3 | 194.2 ± 74.8 | -0.0117 ± 0.0194 | 64 | 6,012 |
| **Western Caribbean** | Belize, Guatemala, Honduras, Mexico | 316,650 | 2220.9 ± 501.9 | 45687.6 ± 13065.1 | 29.8 ± 1.6 | 168.4 ± 64.7 | -0.0220 ± 0.0273 | 20 | 317 |
| **Total** | All listed above | 2,025,295 | 2437.9 ± 488.8 | 36251.8 ± 17791.0 | 29.5 ± 2.4 | 126.7 ± 57.9 | -0.0044 ± 0.0375 | 1,293 | 41,801 |

**Fig. 1.**
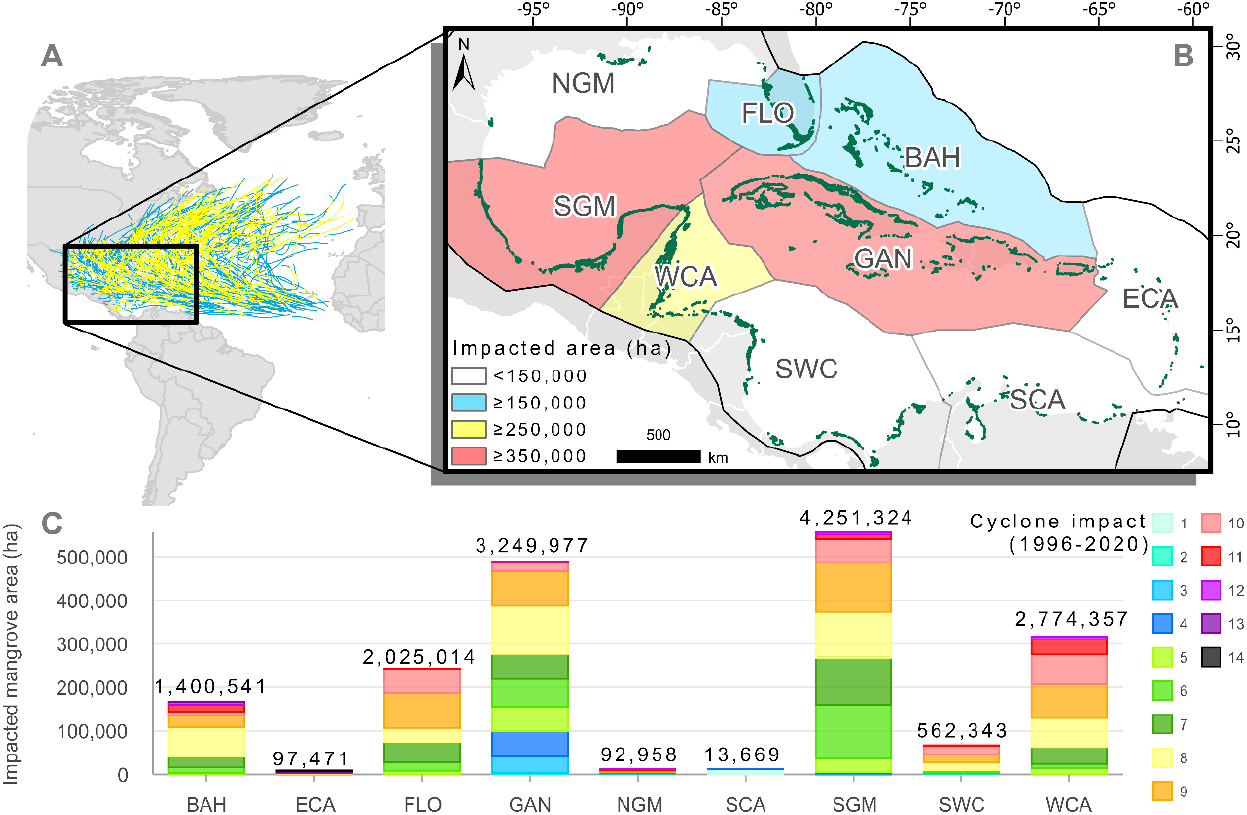
From 368 Atlantic named tropical cyclones over the studied 25 years, the Southern Gulf of Mexico and the Greater Antilles present the largest mangrove impacted area, followed by the Western Caribbean, Floridian, and Bahamian subregions. During this period, some Bahamian and Eastern Caribbean mangroves have suffered more than thirteen cyclones’ landfalls. A) Atlantic named cyclones’ pathways (from storms – in blue to hurricanes – in yellow) from 1996 to 2020, and study site location (black box). B) Zoom-in of the study site showing the distribution of mangroves (in dark green), and its nine subregions colored according to ranges (from < 150,000 to ≥ 350,000 ha) of mangrove area impacted over the last 25 years. C) Cumulative mangrove area impacted by tropical cyclones per subregion (label) with stacked color bars indicating how many hectares have been impacted by cyclones successively (from one to fourteen times) in the last 25 years. BAH = Bahamian subregion, ECA = Eastern Caribbean, FLO = Floridian, GAN = Greater Antilles, NGM = Northern Gulf of Mexico, SCA = Southern Caribbean, SGM = Southern Gulf of Mexico, SWC = Southwestern Caribbean, WCA = Western Caribbean (Spalding et al., 2007). Map lines delineate study areas and do not necessarily depict accepted national boundaries.

The Bahamian, Floridian, and Northern Gulf of Mexico stand out for the highest cumulative wind speeds, while the Southwestern and Western Caribbean subregions stand out for the highest cumulative rainfall from hurricane seasons in the last 40 years (Table 1). The region presents a mean annual maximum temperature of 29.5 °C and a mean annual rainfall of 126.7 mm (CRU et al., 2017), and both temperature and rainfall patterns follow a gradient from higher to lower values from South to North. The region as a whole is observed to be within a period of significant drying (Neelin et al., 2006), with the Bahamian, Greater Antilles, and Western Caribbean subregions presenting the highest drought tendencies (Table 1). The region hosts a total of 1,293 settlements and 41,801 km of roads within 5 km of the coast. While the Northern Gulf of Mexico shows the largest absolute amount of coastal settlements, the Greater Antilles stands out for the largest absolute presence of roads in the coastal zone (Table 1).

### 2.2. Mangrove vulnerability and short-term resilience

We calculated changes in forest greenness through the Normalized Difference Vegetation Index (NDVI) obtained from 30-m resolution, harmonized Landsat-5, −7, and −8 images, to identify damage in impacted mangrove areas, which have been proven reliable to monitor mangrove damage and recovery (Taillie et al., 2020; Lagomasino et al., 2021). Based on ground validated responses (Lagomasino et al., 2021), mangrove damage (vulnerability) followed a threshold change response of −0.2 NDVI (a drop-response of −0.2) between the *ex-ante* NDVI mean value (two years before the disturbance, July 1 to July 1) and the ex-post NDVI mean value (from August 31 to December 31). This 0.2 decline in NDVI following the storm has been reported to show a significant change in canopy cover fraction and canopy height loss (Taillie et al., 2020; Lagomasino et al., 2021) (see example in Appendix S1). To measure post-disturbance recovery (short-term resilience), we focused on the *ex-post* NDVI slope trend for one year after the disturbance (from January 1 to August 31). A negative NDVI trend (or null trend) was classified as loss and a positive trend as recovery. While our short-term losses do not necessarily lead to long-term mangrove mortality, experience in the region showed that 12 months was a reasonable time to expect recovery responses (Lagomasino et al., 2021). Damage and recovery were produced annually at 30-m spatial resolutions for the periods 1996-2020 and 1996-2019, respectively, for the entire NAB region.

We used Global Mangrove Watch data (Bunting et al., 2018) for our mangrove baseline (the year 1996). Tropical cyclones were collected from NOAA’s-IBTrACS (Knapp et al., 2010) and include both tropical storms (wind speed 63-117 km.hr^-1^) and hurricanes (wind speeds ≥ 119 km.h^-1^), following the Saffir-Simpson scale. Impacted areas correspond to predefined buffers of 80-km radio on the linear features of IBTrACS, as plausible areas of major impact. Annual responses of damage and recovery were grouped into one aggregated layer, offering a snapshot vision that substituted time by space. Class point data (i.e., pixel centroids) were randomly extracted from the aggregated layers to assess the drivers for the NAB region and subregions (Spalding et al., 2007). The analysis of the subregions was considered highly relevant due to the large differences in environmental settings (e.g., soil type, mean elevation, geomorphology), frequency of cyclones, socio-economic frameworks, and governance settings among them.

### 2.3. Drivers of mangrove vulnerability and short-term resilience

From an initial database of 77 possible drivers, we applied nonparametric Spearman and Kendall Rank Correlation tests to exclude highly correlated variables, ending with a final pruned-set of 22 possible drivers (see a summary of the variables’ characteristics, and sources in Appendix S2 and correlation matrices in Appendices S3 and S4) as follows:

#### 2.3.1. Weather variables – the year of cyclone impacts

Weather variables (the same year as the cyclone) are from the Copernicus Climate Change Service (C3S) Climate Data Store (CDS) ERA5 reanalysis with 0.25° of spatial resolution and hourly temporal resolution (Hersbach et al., 2018). We produced annual maximum sustained wind speed images from 1996 to 2020 from ERA5’s hourly wind data, which summarizes maximum 3-second wind at 10-m height. Hereafter it is referred to as the ‘wind speed’. We developed two cumulative rainfall products per year, one from May to November (referred to as the ‘hurricane season’) and one for December to April (‘dry season’). For mangroves with repeated cyclone impacts, we estimated mean wind speed and cumulative rainfall values for each pixel. For the ‘undamaged’ class, we took mean values from the 25-year period of analysis.

#### 2.3.2. Climate variables – long-term means and trends

We estimated tropical cyclone recurrence by summing up how many times each pixel intercepted an 80km tropical cyclone buffer over the 25-year period. Five other variables were calculated to capture long-term climate variability in the region: 1) mean annual maximum temperature (°C), 2) the trend of the annual maximum temperature (°C.yr^-1^), 3) mean annual cumulative rainfall (mm), 4) the trend of the annual cumulative precipitation (mm.yr^-1^), and 5) the trend of the annual mean Self-calibrating Palmer Drought Severity Index (scPDSI) (unitless, but a negative trend indicating drought) from 1980 to 2016. For that, we used monthly data from the Climatic Research Unit (CRU et al., 2017) database at 0.5° spatial resolution.

#### 2.3.3. Human infrastructure variables

We generated three layers of distance to human infrastructures, in order to verify whether coastal development may affect mangrove responses to tropical cyclones. These included Euclidean distance at 100 m spatial resolution from croplands, roads, and settlements. The original cropland mask came from the global croplands project, version 1, at 30 m spatial resolution from 2010 (North America; Massey et al., 2017) and 2015 (South America; Zhong et al., 2017). Croplands with an area lower than one hectare were excluded to reduce feature complexity and allow the generation of a single distance raster for the entire region. Road polyline data were taken from the ‘Global Roads Open Access’ dataset, version 1 with accuracies ranging from 530 m to 1,265 m (CIESEN, 2013). The settlement point data we used are from the Global Rural-Urban Mapping Project, version (CIESEN, 2011).

#### 2.3.4. Mangrove structural variables

We included a total of four habitat variables: one for height, and three for canopy cover closeness. We used data on mangrove canopy height from NASA’s ORNL DAAC (Simard et al., 2019), derived from the Shuttle Radar Topography Mission (SRTM) and the RH100 product from the Geoscience Laser Altimeter System (GLAS) instrument aboard ICESat-1. This data is from 2000 at a 30-m resolution.

We included an indicator of canopy cover closure to provide a baseline for pre-cyclonic mangrove conditions. We generated sub-pixel fractional covers of green vegetation, soil, and water from a Spectral Mixture Analysis (SMA). For that, we collected two sets of 100 candidate spectra throughout the region, and across years (and seasons) each dataset corresponded and was applied to Landsat-5 collection 1, tier 1 (from 1994 to 2011), and Landsat-8 collection 1, tier 1 (from 2013 to 2020) (Appendix S5). We ran the images unmixing using Google Earth Engine ‘unmix’ function (Gorelick et al., 2017; Bullock et al., 2020). For mangrove samples with more than one damage, loss/recovery over time, the pre-cyclone fractional cover assigned was associated with the first cyclone impact since 1996.

#### 2.3.5. Geomorphology and soil variables

We selected four geomorphological typologies: Delta, Estuary, Lagoon, and Open Coast, obtained from Worthington et al. (2020). These data were produced using high-resolution coastline layers to map and classify coastal embayment polygons through machine learning. We also generated a 100-m spatial resolution layer of Euclidean distance to the shoreline, using the Prototype Global Shoreline Data (https://shoreline.noaa.gov/data/datasheets/pgs.html). It is based on orthorectified, 2000-era, Landsat imagery and has an accuracy of ca. 50 m.

We include data on soil organic carbon stock at 1-m depth from the global map of mangrove forest soil carbon at 30 m spatial resolution (Sanderman et al., 2018). This database was produced by the 250-m SoilGrid data modeling from various finer resolution explanatory variables such as a digital elevation model, geomorphology map, and vegetation characteristics. The data presents a soil organic density root mean squared error (RMSE) of 10.9 kg/ m^3^.

### 2.4. Statistical analyses and Machine learning classification

We applied descriptive statistics (i.e., median and quartiles per class of vulnerability and recovery), non-parametric statistic tests (Kruskal-Wallis for numerical, and Chi-squared for categorical variables), and machine learning (Random Forest (RF) binary classification) (Breiman, 2001; Kuhn et al., 2020) to characterize and select the most influential drivers of vulnerability and resilience for the NAB region and the subregions. Not all nine subregions hosted enough mangrove data to run RF that required balanced control vs. treatment subsamples. Five subregions, hosting 94% of the total mangrove area in the region, remained for running subregional models: the Bahamian, Floridian, Greater Antilles, Southern Gulf of Mexico, and the Western Caribbean (**Fig. 1**). Thus, twelve models were run: one regional and five subregional models for vulnerability (“damaged vs non-damaged mangroves”) and for post-disturbance resilience (“mangrove recovery vs mangrove loss”) (Appendix S7).

Here, we set 700 trees per model and left the number of predictors at each split to be selected based on the highest overall accuracy. 10-fold cross-validation (10% of data hold-out for validation) tested the accuracy of the models. The relative importance of different drivers relied on the difference between two prediction accuracies on the out-of-bag portion of the data, for each tree, when permuting each predictor variable (Kuhn, 2012). All data analyses were performed using R 4.0.0 (R Core Team, 2020).

## 3. Results

### 3.1. How affected by tropical cyclones are mangroves in the NAB region?

From 1996 to 2020, the NAB region hosted 368 named tropical cyclones (87 are category ≥3, i.e., wind speed ≥178 km.h^-1^) (**Fig. 1A** and **Fig. 1B**). Of the 2 million hectares of mangroves in the NAB originally (Bunting et al., 2018), almost all (93%) were impacted by at least one cyclone landfall. Half of that area (55%) was impacted by eight or more landfalls in the 25-year study period (**Fig. 1C** and Table 2). Geographically, the eastern side of the NAB and Western Caribbean acted as a preferential pathway for cyclones with the Eastern Caribbean subregion hosting the highest number of landfalls: 14 in 25 years (**Fig. 1C** and **Fig. 2A**). These numbers offer an idea of the pressure that mangroves are subjected to with cyclone return intervals (8/25; 14/25). However, the highest absolute areas of impacted mangroves correspond to subregions with larger mangrove extents: the Southern Gulf of Mexico and the Greater Antilles, with ca. 500,000 ha of impacted mangroves each. (**Fig. 1C** and Table 2).

**Table 2.** Original mangrove area in hectare (ha) (1), and which was, *at least once,* impacted by tropical cyclones (2) and damaged (3) from 1996 to 2020, and recovered (4) and lost (5) from 1996 to 2019. Data are presented by subregion and summed for the entire North Atlantic Basin region.

| Subregion | 1. Original area (1996) (ha) | 2. Impact (1996-2020) |  | 3. Damage (1996-2020) |  |  | 4. Recovery (1996-2019) |  |  | 5. Loss (1996-2019) |  |  |
| --- | --- | --- | --- | --- | --- | --- | --- | --- | --- | --- | --- | --- |
|  |  | (ha) | % Impact/Original | (ha) | % Damage/Original | % Damage/Impact | (ha) | % Recovery/Original | % Recovery/Damage | (ha) | % Loss/Original | % Loss/Damage |
| **Bahamian** | 167,475 | 167,475 | 100 | 38,109 | 22.8 | 22.8 | 11,615 | 6.9 | 32.4 | 26,019 | 15.5 | 72.7 |
| **Eastern Caribbean** | 87,945 | 87,945 | 100 | 2,458 | 28.0 | 28.0 | 832 | 9.5 | 45.6 | 964 | 1.1 | 52.9 |
| **Floridian** | 243,409 | 243,409 | 100 | 57,558 | 23.6 | 23.6 | 43,897 | 18.0 | 78.4 | 18,267 | 7.5 | 32.6 |
| **Greater Antilles** | 489,790 | 489,790 | 100 | 104,037 | 21.2 | 21.2 | 54,498 | 11.1 | 53.5 | 45,920 | 9.4 | 45.1 |
| **Northern Gulf of Mexico** | 12,098 | 12,098 | 100 | 662 | 5.5 | 5.5 | 81 | 0.7 | 60.4 | 53 | 0.4 | 39.8 |
| **Southern Caribbean** | 52,552 | 12,428 | 23.6 | 85 | n/a | n/a | 33 | n/a | n/a | 46 | n/a | n/a |
| **Southern Gulf of Mexico** | 559,204 | 556,451 | 99.5 | 38,425 | 6.9 | 6.9 | 16,304 | 2.9 | 50.5 | 18,990 | 3.4 | 58.9 |
| **Southwestern Caribbean** | 175,322 | 67,160 | 38.3 | 8,703 | 5.0 | 13.0 | 4,973 | 2.8 | 63.1 | 3,048 | 1.7 | 38.7 |
| **Western Caribbean** | 316,650 | 316,645 | 100 | 29,877 | 9.4 | 9.4 | 9,560 | 3.0 | 47.2 | 10,578 | 3.3 | 52.2 |
| **Total** | 2,025,295 | 1,874,257 | 92.5 | 279,914 | 13.8 | 14.9 | 141,793 | 7.0 | 55.4 | 123,885 | 6.1 | 48.4 |

**Fig. 2.**
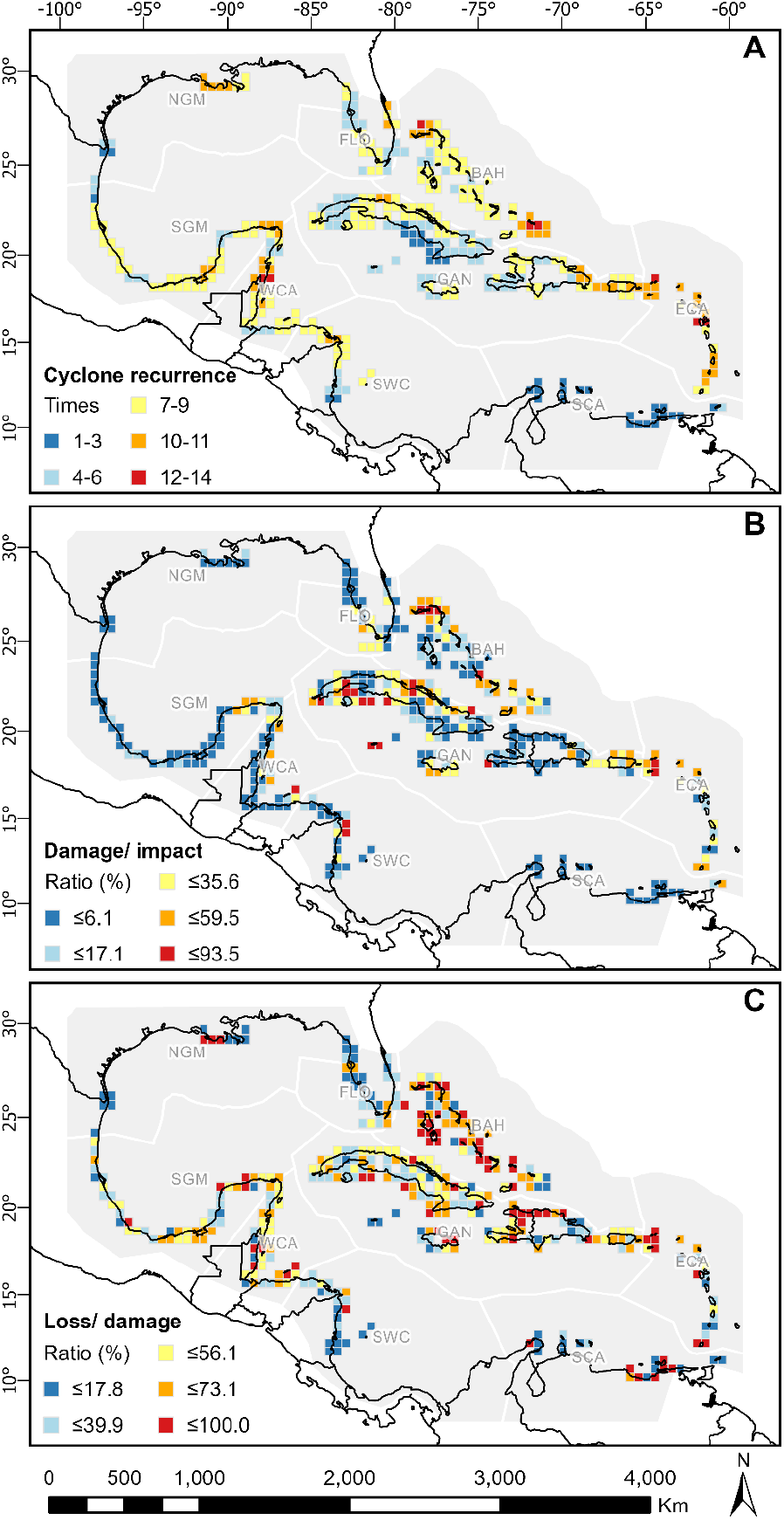
The Eastern side of the Caribbean, mainly, and the Western Caribbean act as preferential pathways of cyclones and, while mangroves see low vulnerability with some hotspots in the central-eastern region, most of them see a high loss of resilience after damage with highlight to the Bahamian (BAH) subregion. Spatial distribution of cyclone recurrence (maximum recurrence at each 0.5-degree cell) (A), damage/ impact (ratio in percentage at each 0.5-degree cell) (B), and loss/ damage (ratio in percentage at each 0.5-degree cell) (C) from 1996 to 2019. BAH = Bahamian subregion, ECA = Eastern Caribbean, FLO = Floridian, GAN = Greater Antilles, NGM = Northern Gulf of Mexico, SCA = Southern Caribbean, SGM = Southern Gulf of Mexico, SWC = Southwestern Caribbean, WCA = Western Caribbean. Map lines delineate study areas and do not necessarily depict accepted national boundaries.

### 3.2. Locating tropical cyclone impacts, mangrove damage, and post-disturbance recovery and loss

While cyclone landings affected almost all mangroves in the NAB region (**Fig. 2A**), the vulnerability was low, suggesting high adaptation to tropical cyclones. Hence, damage represented a small percent of the total area of impacted mangroves, with 14.9% (279,914 ha) of mangroves damaged at least once during our 25-year period of analysis. The central-eastern side of the NAB (Eastern Caribbean, Bahamian, Greater Antilles) concentrated the highest ratios of mangrove damage (damage:impact) with values above 20% (Table 2 and **Fig. 2B**). The Southern Caribbean subregion received little to no impact as this region is located outside of the preferential pathway of the cyclones.

Regional mangrove resilience was, however, rather low with rates of post-disturbance loss (loss:damage) being ca. 48% (Table 2). A total of 123,885 ha mangrove showed no recovery trends in the following 12 months after disturbance. Maximum resilience was observed in the Floridian subregion, with 78.4% of recovery, while the lowest values were within the Bahamian (32.4%) and Eastern Caribbean (45.6%) subregions (Table 2). In absolute numbers, mangrove-rich central-western subregions (i.e., the Greater Antilles, the Western Caribbean, and the Southern Gulf of Mexico) contributed the most to mangrove losses (Table 2 and **Fig. 2C**).

### 3.3. Drivers of mangrove vulnerability

Mangrove vulnerability to tropical cyclones for the period 1996-2020 was mostly influenced by weather properties linked to the cyclonic events: wind speeds and associated high rainfall (leading to flooding) (detailed information on each vulnerability model, accuracies, and descriptive statistics are presented in Appendices S8 to S14). These were common drivers for the NAB region (**Fig. 3A**) and all the subregions (**Fig. 3B-F**). The weight of wind speed and rainfall was most marked in the subregions located on the preferential pathway of cyclones in the NAB region (eastern side of the Caribbean), the Bahamian, Eastern Caribbean, Greater Antilles, and Floridian subregions (**Fig. 3B-D**). Many of these subregions host islands or are Peninsulas, and cyclones pass across them without losing full strength. More continental subregions, such as the Southern Gulf of Mexico or the Western Caribbean saw wind speed and high rainfall as leading drivers of vulnerability, but long-term climate trends, human infrastructure, geomorphology, and pre-cyclone habitat (height and canopy openness), were the most important drivers of mangrove vulnerability to tropical cyclones (**Fig. 3E-F**).

**Fig. 3.**
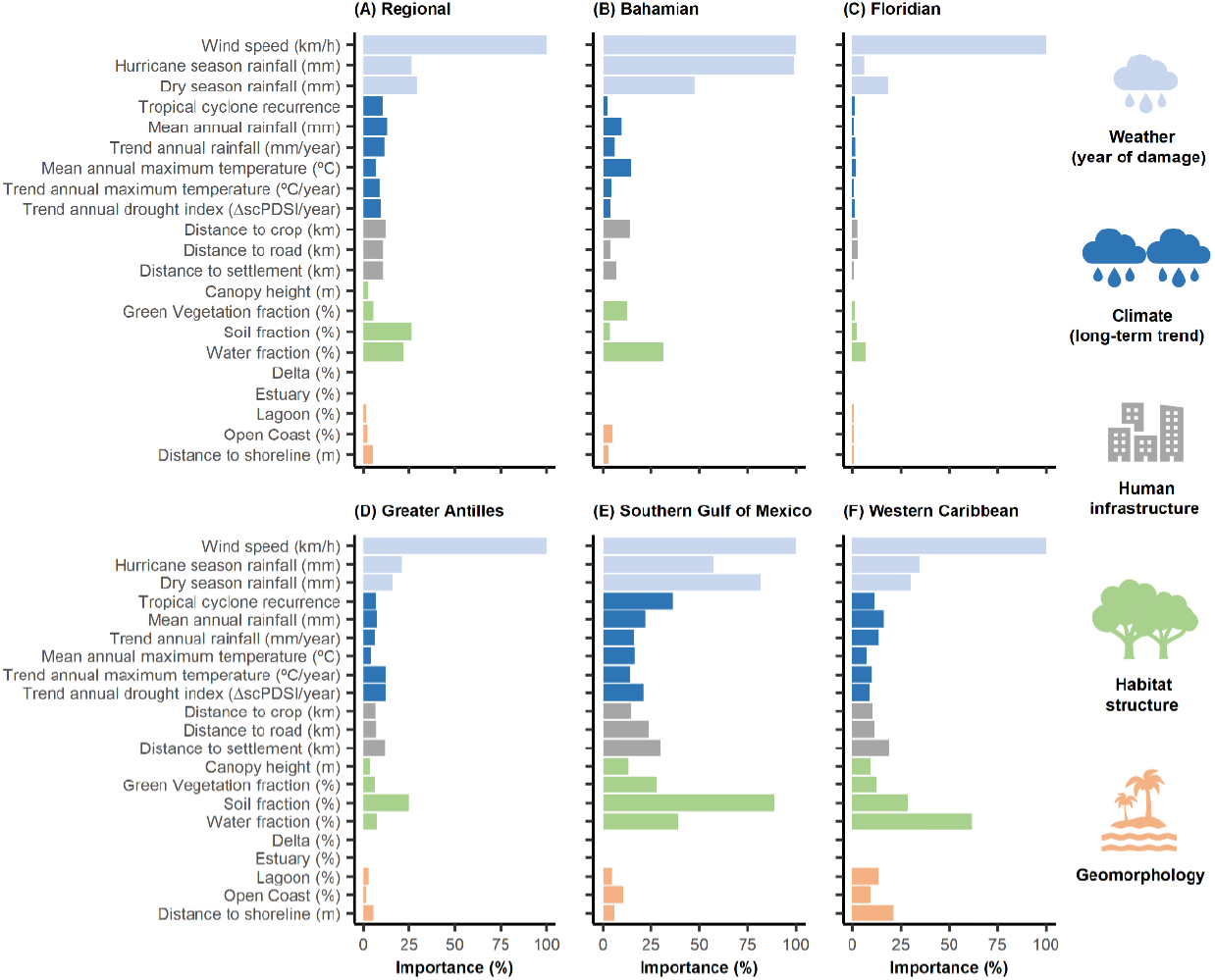
Drivers of mangrove vulnerability at regional (A) and subregional scales (B-E) in the North Atlantic Basin (Gulf of Mexico and Caribbean) suggest cyclonic weather conditions such as wind speeds and associated high levels of rainfall as major drivers of vulnerability in the region. Less insular subregions add other drivers such as pre-cyclone habitat conditions and distance to human infrastructure as extra drivers of mangrove vulnerability. Bars are proportional to the variable importance from mangrove damage vs. non-damage classification trees (n=700). Five out of nine subregions (Spalding et al., 2007) are assessed here (i.e., those comprising ≥ 94% of the mangrove extent in the NAB region: Bahamian, Floridian, Greater Antilles, Western Caribbean, and Southern Gulf of Mexico subregions). Data distributions are statistically different between classes (p-value ≤ 0.05) with the exception of dry season rainfall and distance to settlement in the regional model, and the trend of annual drought index in the Southern Gulf of Mexico and Western Caribbean models.

#### 3.3.1. Thresholds of vulnerability at the regional level

Translated into thresholds for the NAB region, we found that mangrove vulnerability to tropical cyclones was enhanced by insularity with wind speeds ≥107 km.hr^-1^, and high levels of associated cyclonic rainfall (~1050 mm) strongly driving mangrove damage (**Fig. 4 A-B**). Taller mangroves (~10m) were more vulnerable than shorter mangroves (**Fig. 4C**), the same way that lagoonal mangroves were more damaged than open coast, estuaries, and delta mangroves (**Fig. 4D**).

**Fig. 4.**
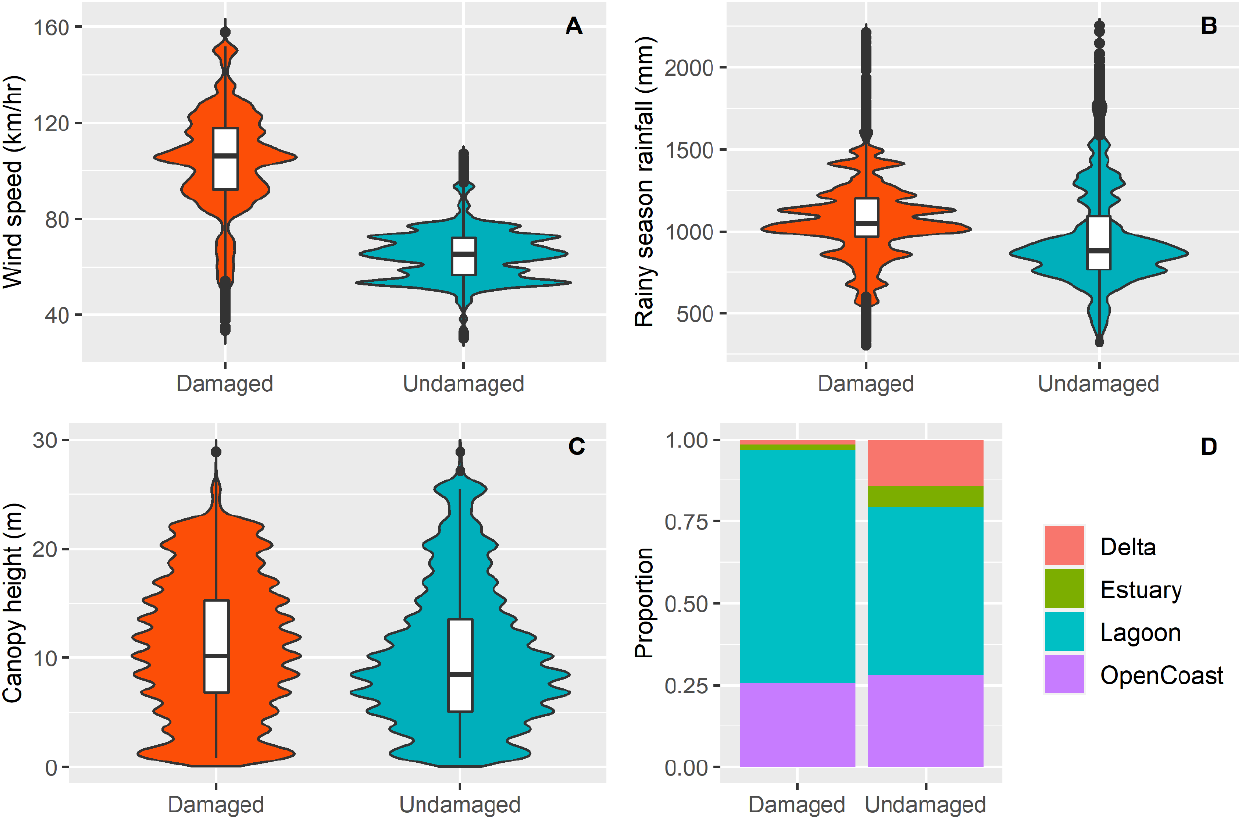
Regional mangrove vulnerability increases where cyclones’ wind speed and rainfall are higher, forests are taller and are in lagoon settings. Violin plots showing mangrove vulnerability (damaged vs undamaged) in the NAB region (Caribbean and Gulf of Mexico) Maximum sustained wind speeds (A), cumulative rainfall in the hurricane (rainy) season (B), canopy height (C), and geomorphological unit (D). The lower and upper edges of the boxes indicate the interval between 25 and 75% of the data distribution, and the central thick line is the median value. Horizontal lines outside the boxes indicate the minimum and maximum values of the dataset that were not outliers, and dots are outliers. Stacked bar plot showing the proportion (0-1) of regional “damaged” and “undamaged” samples per geomorphological setting (D).

#### 3.3.2. Thresholds of vulnerability at the subregional level

On a subregional level, our models showed that wind speed and rainfall interact at multiple timescales to influence mangrove vulnerability (e.g., rainfall during the hurricane season, rainfall preceding the hurricane season, and long-term rainfall trends). The Bahamian subregion was the most vulnerable. There, flooding was as important as wind speed in driving mangrove damage, which was a unique subregional response (**Fig. 3B**). Thus, mangroves in the Bahamian subregion were highly responsive to cumulative rainfall during the hurricane season: 1218 mm vs 763 mm in damaged and undamaged areas, respectively. In this line, mangroves were more vulnerable when shorter (~1.7 m) than when taller (~6.8 m), an exception to the general rule within the NAB. Long-term drought also conditioned the vulnerability of Bahamian mangroves, where damage was more frequent in areas undergoing decreasing rainfall since the ‘80s: median of −0.05 mm.yr^-1^ vs 0.04 mm.yr^-1^ in damaged vs undamaged mangroves (Appendix S10). Rainfall also conditioned vulnerability in other subregions, such as the Southern Gulf of Mexico, where aridity played a role. Thus, mangroves were more damaged in areas with lower median annual rainfall (ca. 77 mm) than higher annual means (118 mm) (Appendix S13).

Human drivers also influenced the vulnerability of mangroves in less insular subregions such as the Southern Gulf of Mexico and the Western Caribbean subregions, where damage was more likely in mangroves two times closer to settlements (16 vs 31 km, damage vs undamaged respectively) (**Fig. 3F** and Appendix S14). Human variables were also key in the Bahamian subregion, where damaged mangroves were up to four-time closer to human infrastructure and to crops than undamaged mangroves in the Bahamian region (**Fig. 3 B** and Supplement).

Geomorphology was very influential in the Greater Antilles subregion where 71% of the damaged mangroves were in lagoonal settings, while 60% of the undamaged mangroves were on the open coast. In the Western Caribbean, distance to the shoreline was important, with mangroves closer to the sea (about 440 m closer) being more likely to be damaged than mangroves farther away (**Fig. 3F**).

### 3.4. Drivers of mangrove post-disturbance resilience

Mangrove short-term resilience for the period 1996-2019 showed a far larger diversity of influences at regional and subregional scales (**Fig 5A** vs. **Fig 3A**), highlighting the importance of local conditions in driving short-term resilience, in contrast to the more uninformed role of weather-related cyclonic properties for vulnerability (detailed information on each recovery model, accuracies, and descriptive statistics are presented in Appendices S15 to S21). Human drivers, habitat pre-cyclone conditions and geomorphology all played influential roles in the NAB and subregional resilience. Regionally, the most remarkable novel influence on mangrove recovery was the yet understudied role of long-term climate trends (e.g., annual cumulative rainfall, Self-Calibrated Palmer Drought Severity Index (scPDSI), and annual maximum temperature). These three variables suggested that mangroves were less resilient to tropical cyclones in sites with drought and heat trends (**Fig. 5A-B**). For example, positive trends in scPDSI (0.03 ΔscPDSI.yr^-1^) were associated with resilience, while areas suffering from drying trends (−0.02 ΔscPDSI. yr^-1^) saw no recovery 12 months after the disturbance. Other drivers of regional resilience also included wind speed, with stronger cyclones (hurricanes cat ≥ 3, wind ≥ 178 km.hr^-1^) leading to higher mangrove recovery. Geomorphology was as important for recovery as it was for vulnerability, with basin mangroves farther from the coast such as lagoonal mangroves being less resilient (Appendix S16).

**Fig. 5.**
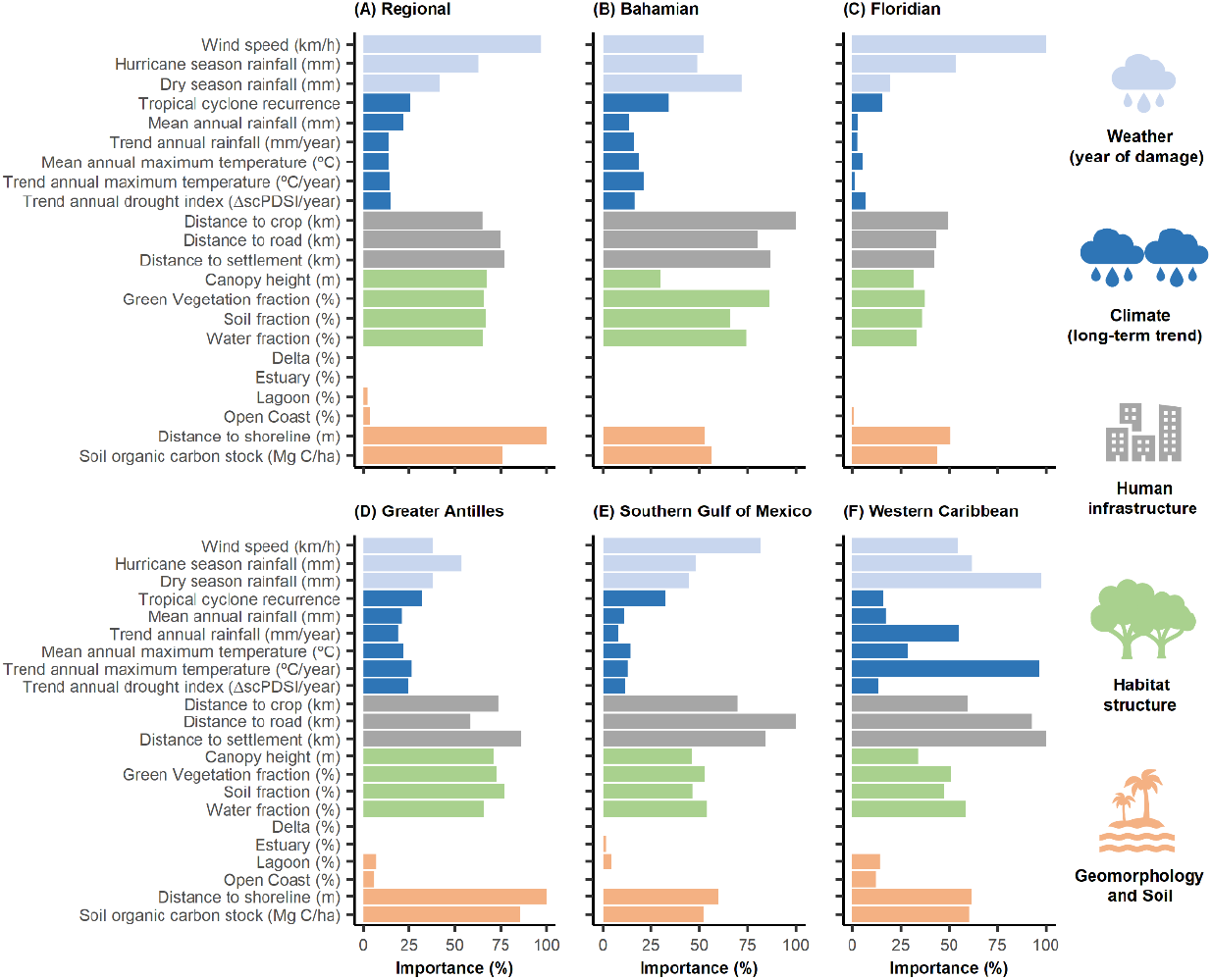
Drivers of mangrove resilience at regional (A) and subregional scales (B-E) in the North Atlantic Basin (Gulf of Mexico and Caribbean) see more diversified site influences, including short and long-term climate impacts, human infrastructure, pre-cyclone habitat condition, geomorphology, distance to shore, and soil carbon. Bars are proportional to the variable importance from mangrove recovery vs. loss classification trees (n=700). Five out of nine subregions (Spalding et al., 2007) are assessed here (i.e., those comprising ≥ 94% of the mangrove extent in the NAB region: Bahamian, Floridian, Greater Antilles, Western Caribbean, and Southern Gulf of Mexico subregions). Data distributions are statistically different between classes (p-value ≤ 0.05) with the exception of distance to road and to shoreline in the regional model, mean annual rainfall and maximum temperature, water fraction and distance to the shoreline in the Bahamian model, cyclone recurrence, soil fraction and distance to the shoreline in the Southern Gulf of Mexico model, and wind speed, cyclone recurrence, distance to settlement, canopy height, green and soil fractions and distance to the shoreline in the Western Caribbean model.

#### 3.4.1. Thresholds of resilience at the regional level

Site conditions played major roles in defining resilience. Mangrove stands with taller trees (13.6 m for recovered mangroves vs 8.5 m for non-recovered), denser canopies (75% of the recovered mangroves had pre-cyclone green vegetation fractions above 80%), and higher soil organic carbon stocks (522 vs 509 Mg C. ha-1 for recovered vs non-recovered, respectively) recover better after disturbance (**Fig. 6C-F**). Distance to shore was an important regional variable influencing mangrove resilience, with closer distances seeing better recovery after cyclones (Appendix S16).

**Fig 6.**
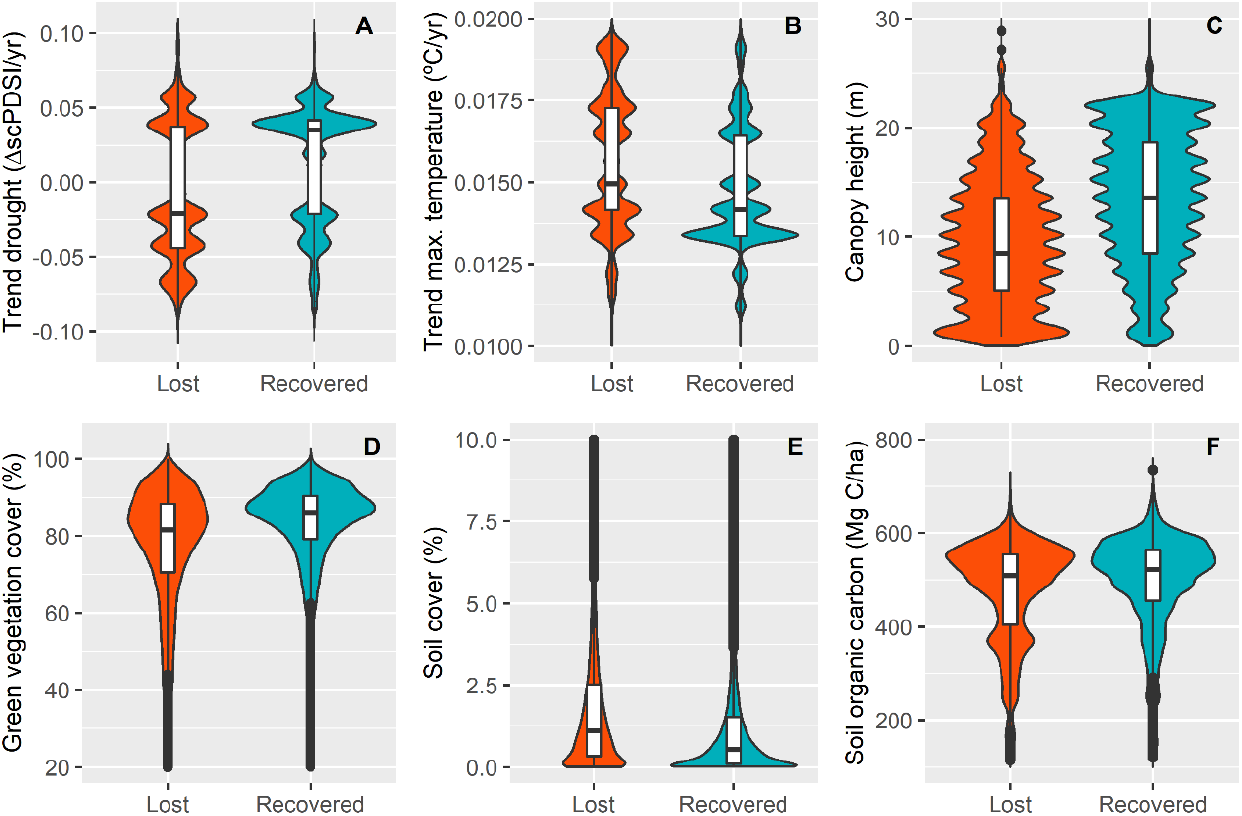
Mangroves in areas suffering from drier and hotter trends, as well as mangroves that are shorter and with lower pre-cyclone canopy cover see less recovery regionally. Soil organic carbon stock seems to promote recovery after a tropical cyclone disturbance. Violin plots showing mangrove ex-post resilience responses (recovered vs lost) in the NAB region. The long-term trend in Self-Calibrated Palmer Drought Severity Index (scPDSI) (A), the long-term trend in annual maximum temperature (B), canopy height (C), pre-cyclone green vegetation cover (D), pre-cyclone soil cover (D), and soil organic carbon stock (E). The lower and upper edges of the boxes indicate the interval between 25 and 75% of the data distribution, and the central thick line is the median value. Horizontal lines outside the boxes indicate the minimum and maximum values of the dataset that were not outliers, and dots are outliers.

#### 3.4.2. Thresholds of resilience at the subregional level

Human infrastructure was a key influencer of mangrove resilience across all subregions (**Fig. 5**). As it was seen for vulnerability, the Bahamian subregion showed clear responses to infrastructure. This was also the case for the Southern Gulf of Mexico and Western Caribbean subregions where human variables displayed more than 80% of the explanatory weight of the recovery model (**Fig. 5B, E-F**). The further the infrastructure was from the mangroves, the better they recovered.

The Western Caribbean subregion captured the best underlying effects of weather conditions in the year of cyclone damage and long-term climate trends on mangrove resilience. Thus, not only did mangroves in drier areas (381 mm) recover worse than in wetter ones (460 mm) but also, areas undergoing long-term drought saw significant post-disturbance losses. With the exception of the Floridian subregion, all the other subregions presented the same pattern of loss of resilience after tropical cyclone disturbances in sites undergoing hotter and drier conditions (**Fig. 5** and Appendices S16-S21). Drying trends in the NAB and its interaction with mangrove resilience is a highlight of this study which requires further research.

As in the regional scale, pre-cyclone habitat conditioned resilience in all the subregions, with higher canopy densities, taller trees, and higher soil organic carbon stock promoting recovery. Exceptions included the Bahamian and the Western Caribbean subregions for tree height, and the Southern Gulf of Mexico for canopy cover (where both recovered and non-recovered mangroves had about 80% of pre-disturbance green cover). Distance to the shoreline was an important variable to model recovery at the regional level, but there was no coherent response among the subregions (**Fig. 5** and Appendices S16-S21).

## 4. Discussion

We highlight that mangrove vulnerability and short-term resilience respond differently to cyclonic impacts in the North Atlantic Basin (NAB) region, which is in line with known trade-offs between ecosystem resistance and resilience to tropical cyclones (Patrick et al., 2022). Mangroves showed low vulnerability (high resistance) to cyclones with only ca. 15% of the vastly impacted area being damaged, but rather low short-term resilience (48% of loss) in areas that were damaged at least once. This is a large percentage for a region where mangroves host numerous post-disturbance traits including rapid re-sprouting, prolific production of seedlings, fast rearrangements in species zonation, and rapid rates of succession and tree growth through their adaptive growth-mortality cycle (Jimenez et al., 1985). Thus, conservation, restoration, and adaptation efforts should focus on those mangroves which are more vulnerable and, mainly, on those less resilient (as highlighted in **Fig. 2**).

There are marked subregional differences between mangrove damage and short-term loss, with east-west and north-south as the main axes. The highest ratios of damage (‘damage:impact’) are mostly concentrated in insular areas along the most preferential pathway of cyclones on the eastern side of the Caribbean: Eastern Caribbean (e.g., Guadeloupe, Grenada, Dominica), Bahamian, and Greater Antilles (e.g., Cuba, Puerto Rico)) subregions. The southern part of the NAB showed less impact, damage, and loss since it is out of the preferential cyclone pathways. When considering the absolute area of damaged mangroves, the western mangrove-rich subregions were also important. Selected damage metrics play, therefore, a key role in prioritizing action. On the other hand, less resilient mangroves, which presented a high ‘loss:damage’ ratio, are in the Bahamian subregion, followed by sites in the Eastern Caribbean, Greater Antilles, Southern Gulf of Mexico, and Western Caribbean mainly.

The characteristics of the cyclones (e.g., wind speed, rainfall) mainly drive vulnerability at the regional level, while short-term resilience is largely driven by site-specific conditions that include long-term climate conditions (e.g., drying trends and drier areas), pre-cyclone habitat conditions and human interventions on the land. The Bahamian subregion stood out for the remarkable vulnerability of its mangroves associated with the hurricane season rainfalls. Its low-lying elevation (e.g., 80% of the landmass is within 1.5 m of mean sea level) and accelerated sea level rise (Kulp & Strauss, 2019; ECLA, 2020) seem to promote impounding conditions that affect mangrove survival. Geomorphology then influences mangroves’ response to cyclone impact, with lagoonal and basin mangroves presenting higher vulnerability and lower short-term resilience regionally. However, it is important to note that geomorphology varies with scale, and its positive or negative impact on mangrove response is associated with other elements such as aridity, karstic environment, and local changes caused by human intervention (Zalivar et al., 2000; Osland et al., 2008; Harris et al., 2010).

Subregions with minimum human intervention close to the mangroves such as the Floridian saw the highest resilience while the Bahamian subregion, with its compounded effects (sea-level rise, urbanization, topography, drought trends) saw the lowest. The Bahamian, Southern Gulf of Mexico, and Western Caribbean subregions are where mangrove vulnerability and resilience are most affected by long-term climate (e.g., declining rainfall) and human development. Both are expected to increase the exposure and vulnerability of ecosystems to extreme events (Sippo et al., 2018), and sites that face both stressors have been reported to exhibit magnified vulnerability in Mexico (Cinco-Castro & Herrera-Silveira, 2020).

The highest mangrove short-term loss in sites showing drought trends is a regional pattern indeed. This is very relevant for the management and restoration of cyclone-damaged mangroves in the NAB since, besides holding six of the driest countries in the world (FAO, 2016), large regions in Central America and the Caribbean are currently undergoing significant drying trends and drying is projected to continue in the future (Neelin et al., 2006). Our results are in line with research on mangrove growth on a global scale, where the influence of precipitation was already highlighted as positively influencing mangrove growth (Simard et al., 2019). Thus, both long-term trends and extreme events of drought represent critical impacts on mangroves, mainly in arid subregions such as the Southern Gulf of Mexico, by increasing site salinity and decreasing mangrove ecophysiological capacity to adapt and be resilient (e.g., Zalivar et al., 2000).

The impacts of increasing hurricanes need to be framed in the context of heavy human influences in the region. Long-term losses of mangroves due to extreme events represented 20% of the total mangrove deforestation in the NAB from 2000 to 2016 (Goldberg et al., 2020). While this contribution is significant and extreme events do play a role in ecosystem stability, the Caribbean has seen massive coastal tourism development in the last 70 years (Pattullo, 2005; Gmelch, 2012) which has long led to mangrove deforestation. These heavy human influences have increased the exposure and vulnerability of coastal communities to these extreme events (Cardona et al., 2012; Ramenzoni et al., 2020) and act as amplifiers of cyclone risks. Interestingly, distance to human infrastructure lost its relevance in the Floridian mangroves. Even though this subregion has a high presence of settlements in the coastal zone, most of its mangroves are under protected status within the US Everglades National Park, which might be what makes them an exception in the region.

Living shorelines (mangroves, reefs, seagrasses, coastal marshes) provide more effective protection against cyclones and better facilitate post-disturbance socio-economic recovery than hard shorelines (Smith et al., 2018; Hochard et al., 2019; Menéndez et al., 2020; Zhu et al., 2020). While the region has multiple legal instruments to protect the coastal and marine ecosystems, donors have not yet fostered the value of NbS (i.e., Green and Green-gray Infrastructures) in their regional budgets for post-disaster reconstruction, where mangrove conservation and restoration seem to remain un-earmarked. Placing ecosystem health - and not only human safety - as part of regional risk reduction strategies, is an urgent task in the region.

Mangrove and socio-economic resilience in the NAB are predicted to worsen. The observed increase in frequency and intensity of cyclones (Bacmeister et al., 2018; Emanuel, 2021; Wang & Toumi, 2021) – here, we show a mean rate of eight cyclones’ impact within the 25 years of study – is at odds with recovery times: 1) the reported mangrove recovery times of 20 years (Lugo, 1980; Jimenez et al., 1985; Roth, 1992; Alongi, 2008; Krauss & Osland; 2020) and 2) the economic recovery time after extreme events in small economies such as the Antillean Islands (ca. 30 years) (Hsiang & Jina, 2014; López-Calva, 2019). In addition, the compounded effects of simultaneous extreme events (ENSO droughts + hurricanes) and underlying climate trends (long-term drying trend in the region + hurricanes) are expected to enhance mortality in a yet not well understood but observable manner, such as the abnormal mangrove mortality registered in 2017 (Taillie et al., 2020). Managers will therefore need to focus on enhancing mangrove adaptation to compound effects of climate change and minimizing human drivers that can make mangroves more vulnerable and reduce their resilience after cyclone impacts (e.g., fragmentation, eutrophication, distance to roads, croplands, settlements).

Our findings have direct and profound implications for ecosystem conservation and coastal protection that interest a broad community of scientists and practitioners. There is an urgent need to reduce human impacts on mangroves and promote ecosystem functionality restoration, following site-specific pressures and needs, as part of risk reduction policies and as part of reconstruction funds after mega-hurricane seasons. We particularly aim to raise awareness among coastal managers and donors to value the role of mangroves restoration and conservation as fundamental NbS against the increasing cyclonic activity in the NAB region.

## Supporting information

Supplement_Amaraletal_Drivers

## Acknowledgments

This research was made possible thanks to the generous support from the BNP-PARIBAS Foundation on their 2019 Climate and Biodiversity Initiative call, through the CORESCAM (“Coastal Biodiversity Resilience to Increasing Extreme Events in Central America”) project. We thank Dr. Kamoru A. Lawal and Prof Mark New for providing long-term climate data and Prof. Helio Garcia Leite for assistance with the statistical analysis. The Smithsonian Marine Station contribution.

## Notes

### Competing Interest Statement

The authors have declared no competing interest.

