## Supplement_Amaraletal_Drivers for "Drivers of mangrove vulnerability and resilience to tropical cyclones in the North Atlantic Basin"

**Supporting information** for **"**Drivers of mangrove vulnerability and resilience to tropical cyclones in the North Atlantic Basin**"**

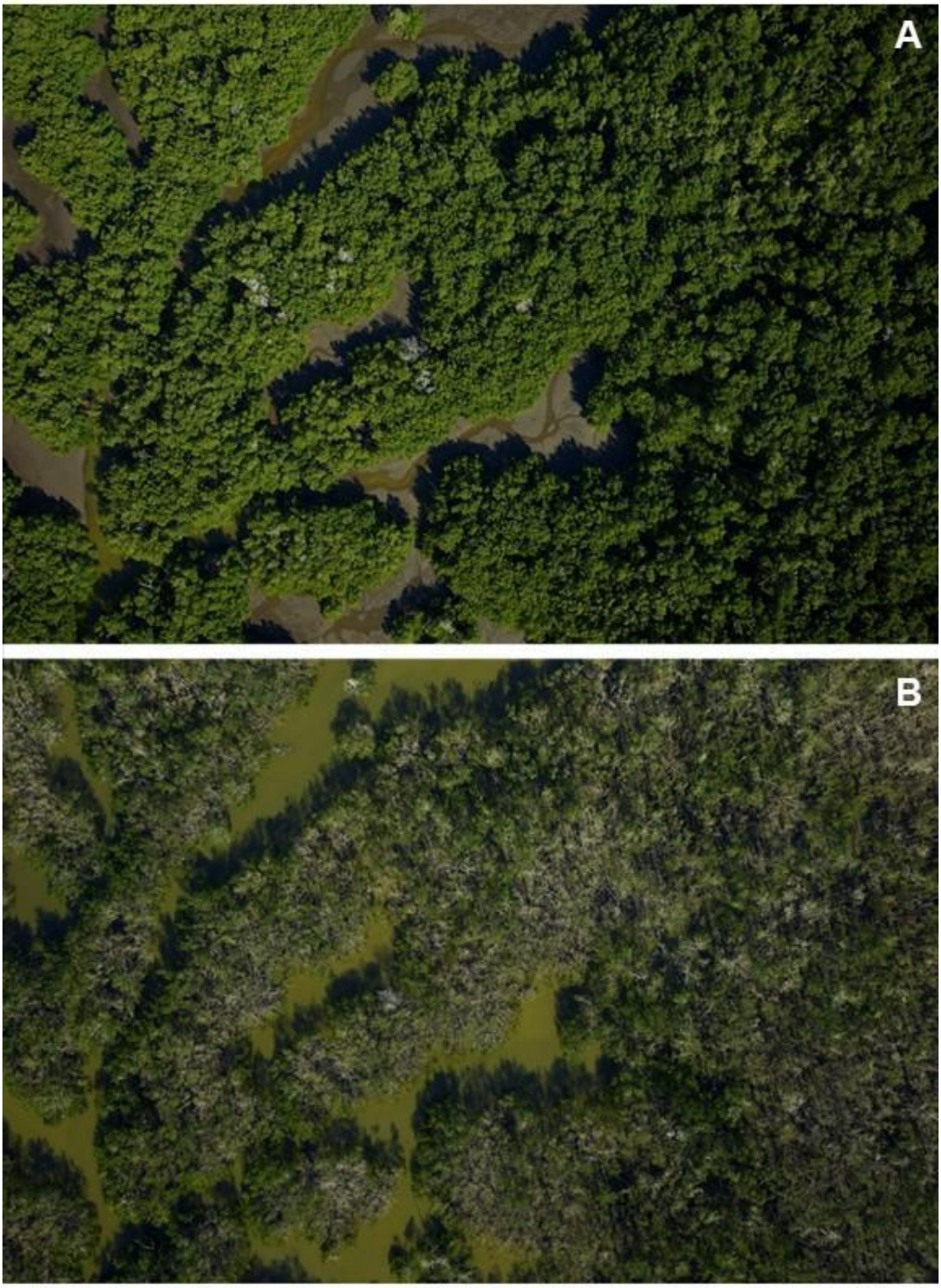


**Appendix S1.** Example of hurricane-driven canopy damage. NASA's G-LiHT Aerial Photography obtained over Floridian mangrove before (A) and after (B) the 2017 hurricane season (available at: https://gliht.gsfc.nasa.gov/index.php?section=49).

**Appendix S2.** Description of the original spatial datasets used in the study.

| Variable | Original data (format) | Spatial resolution | Temporal resolution | Time range | Source | Access |
|---|---|---|---|---|---|---|
| *Wind speed (km/hour)* | *10m wind gust since previous post-processing (pixel)* | 0.25 degree | hourly | 1996-2020 | Copernicus Climate Change Service (C3S) Climate Data Store (CDS) | https://cds.climate.copernicus.eu/cdsapp#!/dataset/reanalysis-era5-single-levels?tab=form |
| *Rainy season rainfall (mm)* | *convective precipitation (pixel)* | 0.25 degree | hourly | 1996-2020 | Copernicus Climate Change Service (C3S) Climate Data Store (CDS) | https://cds.climate.copernicus.eu/cdsapp#!/dataset/reanalysis-era5-single-levels?tab=form |
| *Dry season rainfall (mm)* | *convective precipitation (pixel)* | 0.25 degree | hourly | 1996-2020 | Copernicus Climate Change Service (C3S) Climate Data Store (CDS) | https://cds.climate.copernicus.eu/cdsapp#!/dataset/reanalysis-era5-single-levels?tab=form |
| *Tropical cyclone recurrence* | *tropical cyclone track (polyline)* | 0.10 degree | episodic | 1996-2020 | National Oceanic and Atmospheric Administration (NOAA) | https://www.ncdc.noaa.gov/ibtracs/index.php?name=ib-v4-access |
| *Mean annual rainfall (mm)* | *observed precipitation (pixel)* | 0.50 degree | monthly | 1982-2016 | Climatic Research Unit (CRU) | http://data.ceda.ac.uk/badc/cru/data/cru_ts/cru_ts_3.25/ |
| *Trend rainfall (mm/year)* | *observed precipitation (pixel)* | 0.50 degree | monthly | 1982-2016 | Climatic Research Unit (CRU) | http://data.ceda.ac.uk/badc/cru/data/cru_ts/cru_ts_3.25/ |
| *Mean annual maximum temperature (ºC)* | *monthly average daily maximum temperature (pixel)* | 0.50 degree | monthly | 1982-2016 | Climatic Research Unit (CRU) | http://data.ceda.ac.uk/badc/cru/data/cru_ts/cru_ts_3.25/ |
| *Trend maximum temperature (ºC/year)* | *monthly average daily maximum temperature (pixel)* | 0.50 degree | monthly | 1982-2016 | Climatic Research Unit (CRU) | http://data.ceda.ac.uk/badc/cru/data/cru_ts/cru_ts_3.25/ |
| *Trend drought index (ΔscPDSI/year)* | *self-calibrating Palmer Drought Severity Index (scPDSI) (pixel)* | 0.50 degree | monthly | 1982-2016 | Osborn et al. (2017) | https://crudata.uea.ac.uk/cru/data/drought/ |
| *Distance to cropland (km)* | *cropland (polygon)* | 1 arc second | none | 2010 and 2015 | Massey et al. (2017) and Zhong et al. (2017) | https://lpdaac.usgs.gov/news/release-of-gfsad-30-meter-cropland-extent-products/ |
| *Distance to road (km)* | *road (polyline)* | 4.2 arc second | none | 1980-2010 | Socioeconomic Data and Applications Center (SEDAC) | https://sedac.ciesin.columbia.edu/data/set/groads-global-roads-open-access-v1 |

It continues…

**Appendix S2.** continuation.

| Variable | Original data (format) | Spatial resolution | Temporal resolution | Time range | Source | Access |
|---|---|---|---|---|---|---|
| *Distance to settlement (km)* | *settlement (point)* | 30 arc second | none | 1990-2000 | Socioeconomic Data and Applications Center (SEDAC) | https://sedac.ciesin.columbia.edu/data/set/grump-v1-settlement-points-rev01 |
| *Canopy height (m)* | *canopy height (pixel)* | 1 arc second | none | 2000-2009 | Simard et al. (2019) | https://daac.ornl.gov/cgi-bin/dsviewer.pl?ds_id=1665 |
| *Green vegetation fraction (%)* | *surface reflectance image (pixel)* | 1 arc second | biannual | 1994-2020 | Present study | https://github.com/CibeleAmaral/ClimBiodiv |
| *Soil fraction (%)* | *surface reflectance image (pixel)* | 1 arc second | biannual | 1994-2020 | Present study | https://github.com/CibeleAmaral/ClimBiodiv |
| *Water fraction (%)* | *surface reflectance image (pixel)* | 1 arc second | biannual | 1994-2020 | Present study | https://github.com/CibeleAmaral/ClimBiodiv |
| *Delta (%)* | *delta (polygon)* | 0.8 arc second | none | 1996-2016 | Worthington et al. (2020) | https://data.unep-wcmc.org/datasets/48 |
| *Estuary (%)* | *estuary (polygon)* | 0.8 arc second | none | 1996-2016 | Worthington et al. (2020) | https://data.unep-wcmc.org/datasets/48 |
| *Lagoon (%)* | *lagoon (polygon)* | 0.8 arc second | none | 1996-2016 | Worthington et al. (2020) | https://data.unep-wcmc.org/datasets/48 |
| *Open Coast (%)* | *open coast (polygon)* | 0.8 arc second | none | 1996-2016 | Worthington et al. (2020) | https://data.unep-wcmc.org/datasets/48 |
| *Distance to shoreline (m)* | *shoreline (polyline)* | 1 arc second | none | 2000-present | National Oceanic and Atmospheric Administration (NOAA) | https://shoreline.noaa.gov/data/datasheets/pgs.html |
| *Soil organic carbon stock (Mg C/ha)* | *organic carbon stock (pixel)* | 1 arc second | none | undefined | Sanderman et al. (2018) | https://dataverse.harvard.edu/dataset.xhtml?persistentId=doi:10.7910/DVN/OCYUIT |

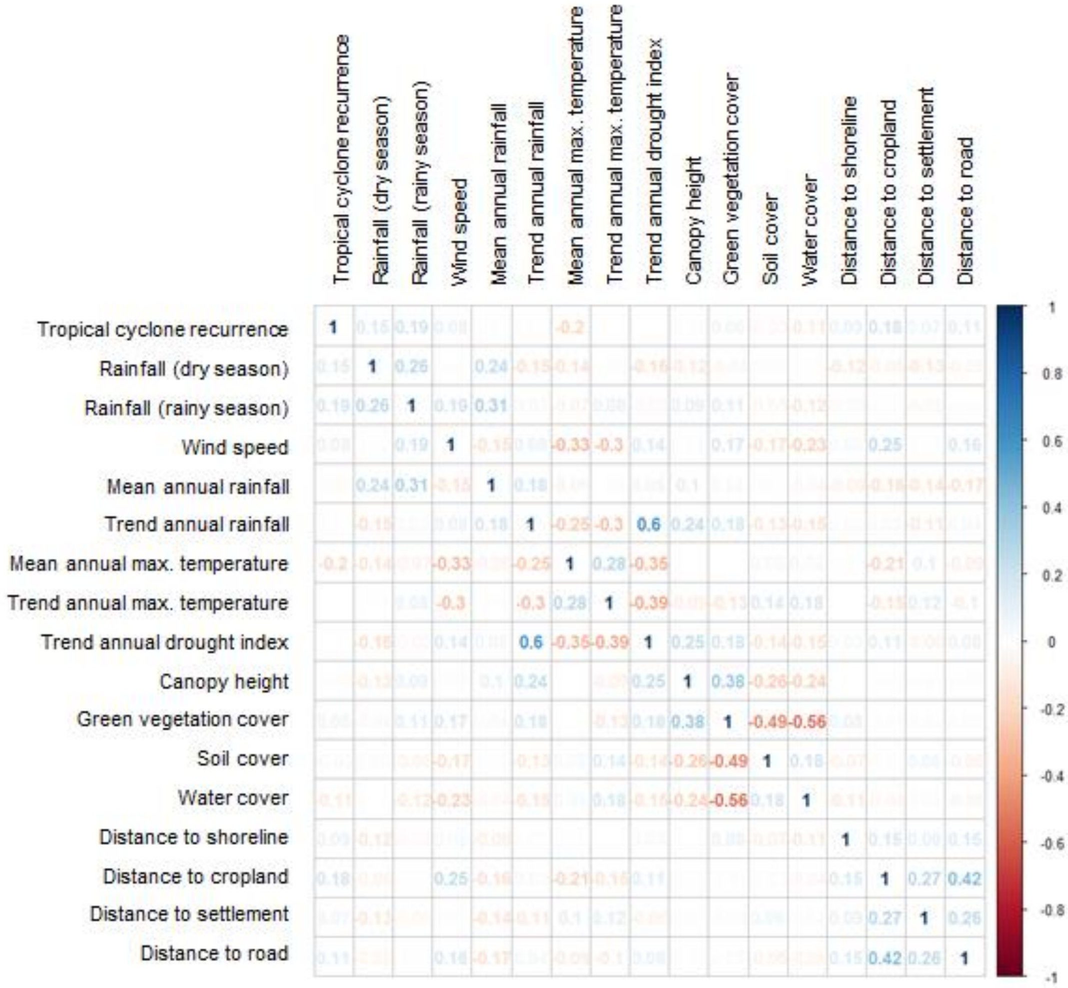


**Appendix S3.** Kendall rank correlation (τ) matrix between numeric variables from the vulnerability model dataset (n = 105,370).

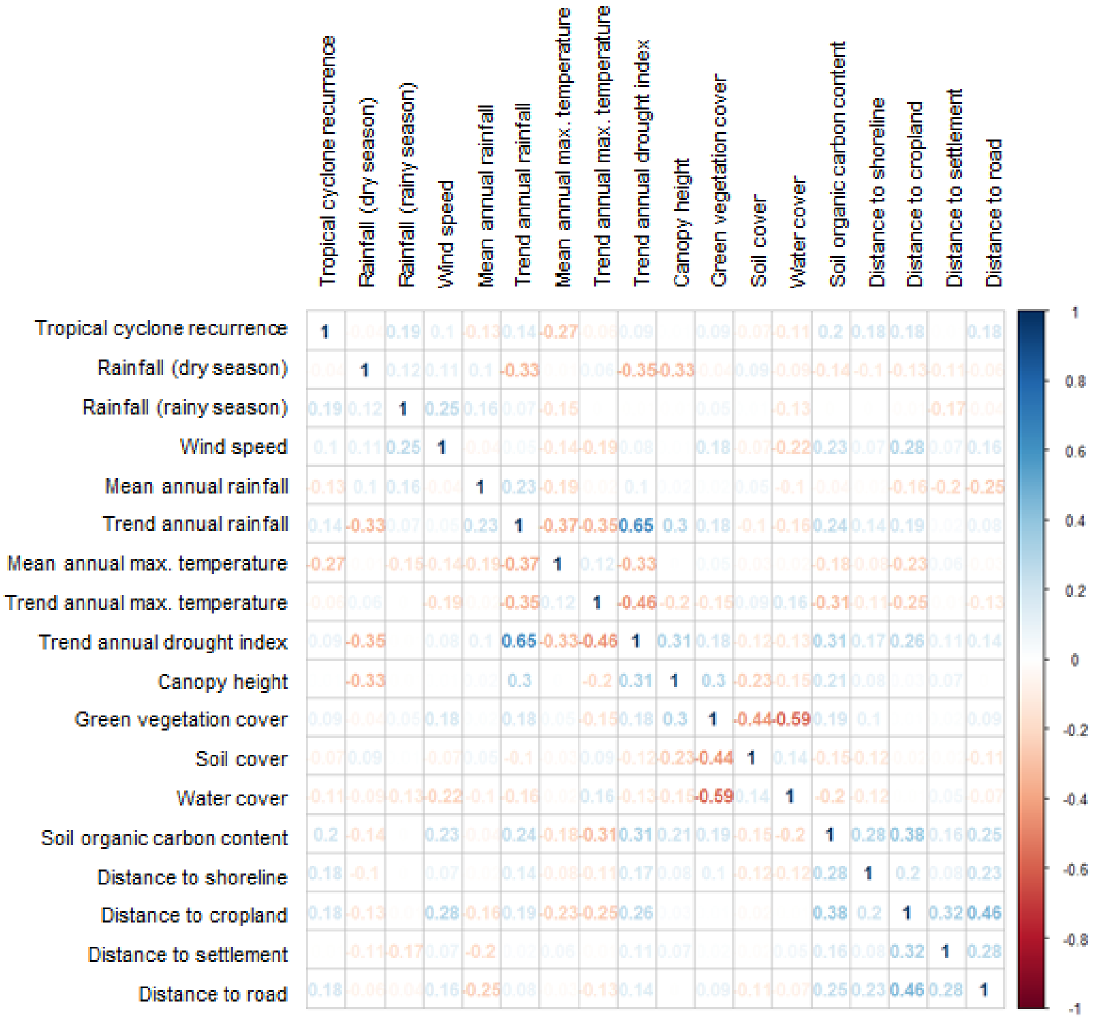


**Appendix S4.** Kendall rank correlation (τ) matrix between numeric variables from the recovery model dataset (n = 118,431).

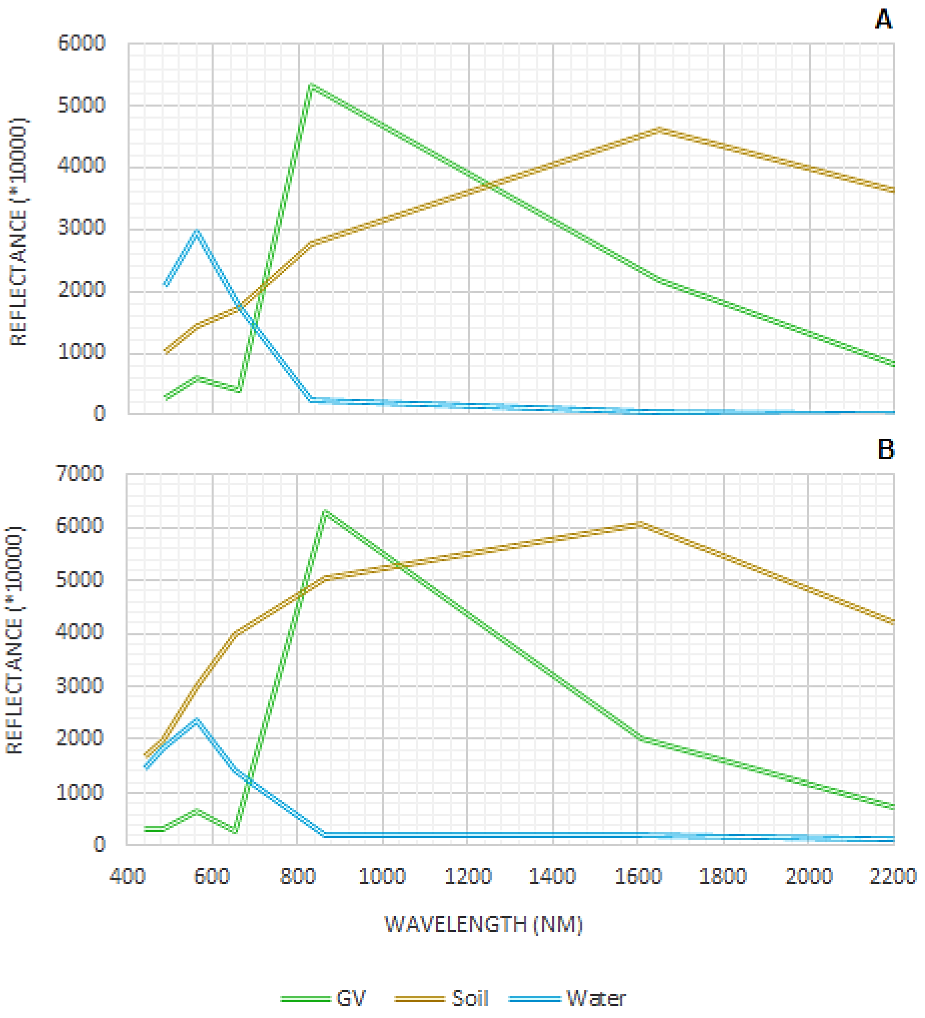


**Appendix S5.** Green vegetation (GV), soil, and water endmembers selected for unmixing Thematic Mapper (Landsat 5) images (**A**) and Operational Land Imager (Landsat 8) images (**B**) from the Caribbean and Gulf of Mexico region.

**Appendix S7**. Sample size for random forest classification by mangrove response (i.e., vulnerability and recovery) model, and subregion.

| Model | Vulnerability | | | Recovery | | |
|---|---|---|---|---|---|---|
| | Damaged | Undamaged | Total | Lost | Recovered | Total |
| **Regional** | 53000 | 52370 | 105370 | 59109 | 59322 | 118431 |
| **Bahamian** | 4200 | 4151 | 8351 | 2794 | 2725 | 5519 |
| **Floridian** | 9900 | 9885 | 19785 | 18112 | 18017 | 36129 |
| **Greater Antilles** | 15900 | 15813 | 31713 | 21095 | 21033 | 42128 |
| **Southern Gulf of Mexico** | 5704 | 5800 | 11504 | 5387 | 5352 | 10739 |
| **Western Caribbean** | 443 | 450 | 893 | 235 | 221 | 456 |

**Appendix S8.** The best number of predictors at each split in tree classification (mtry), and related overall accuracy (%) of each vulnerability random forest model.

| Vulnerability model | mtry | overall accuracy (%) |
|---|---|---|
| **Regional** | 9 | 99.75 |
| **Bahamian** | 9 | 99.71 |
| **Floridian** | 9 | 99.97 |
| **Greater Antilles** | 15 | 99.74 |
| **Southern Gulf of Mexico** | 9 | 99.37 |
| **Western Caribbean** | 9 | 97.50 |

**Appendix S9.** Vulnerability model data distribution from the entire North Atlantic Basin region. Median [first; and third quartiles], and p-value from Kruskal-Wallis median test for each numeric variable by class. For Geomorphology classes, count of samples (and frequency, in percentage), and p-value from Chi-squared test.

| | **Damaged** *N=53000* | **Undamaged** *N=52370* | **p-value** |
|---|---|---|---|
| Tropical cyclone recurrence | 8.00 [7.00;9.00] | 8.00 [6.00;9.00] | <0.001 |
| Rainfall (dry season) (mm) | 162 [131;216] | 173 [118;218] | 0.166 |
| Rainfall (rainy season) (mm) | 1050 [964;1205] | 881 [767;1109] | 0.000 |
| Wind speed (km/h) | 106 [92.3;118] | 65.2 [56.4;71.9] | 0.000 |
| Mean annual rainfall (mm) | 105 [100;116] | 111 [102;131] | 0.000 |
| Trend annual rainfall (mm/yr.) | 0.23 [0.15;0.33] | 0.20 [0.09;0.38] | <0.001 |
| Mean annual max. temperature (ºC) | 29.4 [28.6;30.2] | 30.1 [29.2;31.6] | 0.000 |
| Trend annual max. temperature (ºC/yr.) | 0.01 [0.01;0.02] | 0.02 [0.01;0.03] | 0.000 |
| Trend drought index (∆scPDSI/yr.) | 0.00 [-0.04;0.04] | -0.02 [-0.05;0.03] | 0.000 |
| Canopy height (m) | 10.2 [6.79;15.3] | 8.48 [5.09;13.6] | <0.001 |
| Green vegetation fractional cover | 0.83 [0.74;0.89] | 0.77 [0.67;0.83] | 0.000 |
| Soil fractional cover | 0.01 [0.00;0.02] | 0.02 [0.01;0.05] | 0.000 |
| Water fractional cover | 0.07 [0.04;0.10] | 0.09 [0.08;0.11] | 0.000 |
| Geomorphology class: | | | 0.000 |
| Delta | 828 (1.56%) | 7512 (14.3%) | |
| Estuary | 977 (1.84%) | 3226 (6.16%) | |
| Lagoon | 37601 (70.9%) | 26925 (51.4%) | |
| OpenCoast | 13594 (25.6%) | 14707 (28.1%) | |
| Distance to shoreline (m) | 671 [224;1749] | 539 [141;1513] | <0.001 |
| Distance to cropland (m) | 12148 [2283;48071] | 4627 [1640;17709] | 0.000 |
| Distance to settlement (m) | 28103 [14988;43063] | 27272 [14407;44284] | 0.136 |
| Distance to road (m) | 9247 [4319;19084] | 6760 [3220;13149] | 0.000 |

**Appendix S10.** Vulnerability model data distribution from the Bahamian sub-region. Median [first; and third quartiles], and p-value from Kruskal-Wallis median test for each numeric variable by class. For Geomorphology classes, count of samples (and frequency, in percentage), and p-value from Chi-squared test.

| | **Damaged** *N=4200* | **Undamaged** *N=4151* | **p-value** |
|---|---|---|---|
| Tropical cyclone recurrence | 9.00 [8.00;10.0] | 8.00 [8.00;8.00] | <0.001 |
| Rainfall (dry season) (mm) | 301 [288;370] | 182 [172;242] | 0.000 |
| Rainfall (rainy season) (mm) | 1218 [1124;1420] | 763 [755;861] | 0.000 |
| Wind speed (km/h) | 110 [92.5;115] | 66.3 [65.2;87.2] | 0.000 |
| Mean annual rainfall (mm) | 105 [104;105] | 102 [99.2;110] | <0.001 |
| Trend annual rainfall (mm/yr.) | -0.05 [-0.10;0.04] | 0.04 [-0.06;0.05] | <0.001 |
| Mean annual max. temperature (ºC) | 28.5 [28.5;28.5] | 28.8 [28.8;29.9] | 0.000 |
| Trend annual max. temperature (ºC/yr.) | 0.02 [0.02;0.02] | 0.02 [0.02;0.02] | <0.001 |
| Trend drought index (∆scPDSI/yr.) | -0.07 [-0.07;-0.07] | -0.06 [-0.06;-0.06] | 0.000 |
| Canopy height (m) | 1.70 [0.85;3.39] | 6.79 [3.39;8.48] | 0.000 |
| Green vegetation fractional cover | 0.54 [0.43;0.66] | 0.46 [0.38;0.54] | <0.001 |
| Soil fractional cover | 0.02 [0.01;0.05] | 0.17 [0.07;0.24] | 0.000 |
| Water fractional cover | 0.15 [0.09;0.23] | 0.12 [0.09;0.17] | <0.001 |
| Geomorphology class: | | | 0.000 |
| Lagoon | 134 (3.19%) | 1586 (38.2%) | |
| OpenCoast | 4066 (96.8%) | 2565 (61.8%) | |
| Distance to shoreline (m) | 224 [100;1500] | 424 [141;1265] | <0.001 |
| Distance to cropland (m) | 13062 [8768;39046] | 54818 [30464;83579] | <0.001 |
| Distance to settlement (m) | 22811 [13724;27890] | 52212 [30288;64563] | 0.000 |
| Distance to road (m) | 8431 [5132;12434] | 23889 [9903;34712] | 0.000 |

**Appendix S11.** Vulnerability model data distribution from the Floridian sub-region. Median [first; and third quartiles], and p-value from Kruskal-Wallis median test for each numeric variable by class. For Geomorphology classes, count of samples (and frequency, in percentage), and p-value from Chi-squared test.

| | **Damaged** | **Undamaged** | **p-value** |
|---|---|---|---|
| | ***N=9900*** | ***N=9885*** | |
| Tropical cyclone recurrence | 9.00 [8.00;9.00] | 9.00 [7.00;9.00] | <0.001 |
| Rainfall (dry season) (mm) | 134 [128;162] | 169 [151;195] | 0.000 |
| Rainfall (rainy season) (mm) | 1043 [997;1225] | 863 [841;893] | 0.000 |
| Wind speed (km/h) | 106 [101;127] | 72.7 [69.2;76.7] | 0.000 |
| Mean annual rainfall (mm) | 102 [100;112] | 112 [103;116] | <0.001 |
| Trend annual rainfall (mm/yr.) | 0.33 [0.32;0.51] | 0.51 [0.32;0.55] | <0.001 |
| Mean annual max. temperature (ºC) | 28.8 [28.6;29.2] | 28.6 [28.6;29.0] | <0.001 |
| Trend annual max. temperature (ºC/yr.) | 0.01 [0.01;0.01] | 0.01 [0.01;0.01] | <0.001 |
| Trend drought index (∆scPDSI/yr.) | 0.04 [0.04;0.04] | 0.04 [0.03;0.06] | 0.003 |
| Canopy height (m) | 13.6 [10.2;18.7] | 10.2 [6.79;13.6] | 0.000 |
| Green vegetation fractional cover | 0.85 [0.80;0.89] | 0.77 [0.69;0.82] | 0.000 |
| Soil fractional cover | 0.01 [0.00;0.02] | 0.02 [0.01;0.05] | 0.000 |
| Water fractional cover | 0.07 [0.05;0.09] | 0.09 [0.08;0.11] | 0.000 |
| Geomorphology class: | | | <0.001 |
| Estuary | 124 (1.25%) | 884 (8.94%) | |
| Lagoon | 9036 (91.3%) | 7885 (79.8%) | |
| OpenCoast | 740 (7.47%) | 1116 (11.3%) | |
| Distance to shoreline (m) | 1105 [400;2500] | 500 [141;1300] | <0.001 |
| Distance to cropland (m) | 53868 [35878;61576] | 30771 [8220;50741] | 0.000 |
| Distance to settlement (m) | 40436 [12784;48897] | 14217 [5536;43064] | <0.001 |
| Distance to road (m) | 14947 [5730;30099] | 8201 [3500;20462] | <0.001 |

**Appendix S12.** Vulnerability model data distribution from the Greater Antilles sub-region. Median [first; and third quartiles], and p-value from Kruskal-Wallis median test for each numeric variable by class. For Geomorphology classes, count of samples (and frequency, in percentage), and p-value from Chi-squared test.

| | **Damaged** *N=15900* | **Undamaged** *N=15813* | **p-value** |
|---|---|---|---|
| Tropical cyclone recurrence | 8.00 [6.00;9.00] | 7.00 [5.00;8.00] | 0.000 |
| Rainfall (dry season) (mm) | 191 [158;216] | 130 [96.8;209] | 0.000 |
| Rainfall (rainy season) (mm) | 1015 [870;1156] | 837 [755;919] | 0.000 |
| Wind speed (km/h) | 107 [93.8;118] | 69.5 [63.8;73.1] | 0.000 |
| Mean annual rainfall (mm) | 108 [104;121] | 110 [104;122] | <0.001 |
| Trend annual rainfall (mm/yr.) | 0.20 [0.16;0.26] | 0.24 [0.17;0.30] | 0.000 |
| Mean annual max. temperature (ºC) | 30.1 [29.8;30.5] | 30.2 [30.0;30.7] | <0.001 |
| Trend annual max. temperature (ºC/yr.) | 0.02 [0.01;0.02] | 0.01 [0.01;0.02] | <0.001 |
| Trend drought index (∆scPDSI/yr.) | -0.02 [-0.04;0.02] | -0.01 [-0.02;0.03] | <0.001 |
| Canopy height (m) | 10.2 [6.79;15.3] | 8.48 [5.09;13.6] | <0.001 |
| Green vegetation fractional cover | 0.86 [0.77;0.92] | 0.78 [0.70;0.82] | 0.000 |
| Soil fractional cover | 0.01 [0.00;0.02] | 0.01 [0.00;0.03] | <0.001 |
| Water fractional cover | 0.05 [0.02;0.08] | 0.09 [0.08;0.11] | 0.000 |
| Geomorphology class: | | | 0.000 |
| Delta | 21 (0.13%) | 947 (5.99%) | |
| Estuary | 32 (0.20%) | 439 (2.78%) | |
| Lagoon | 11324 (71.2%) | 4890 (30.9%) | |
| OpenCoast | 4523 (28.4%) | 9537 (60.3%) | |
| Distance to shoreline (m) | 608 [224;1421] | 500 [200;1140] | <0.001 |
| Distance to cropland (m) | 2915 [1208;12702] | 2435 [1077;5590] | <0.001 |
| Distance to settlement (m) | 24253 [15608;34358] | 21132 [14490;33381] | <0.001 |
| Distance to road (m) | 7845 [3614;15593] | 6389 [3202;11314] | <0.001 |

**Appendix S13.** Vulnerability model data distribution from the Southern Gulf of Mexico sub-region. Median [first; and third quartiles], and p-value from Kruskal-Wallis median test for each numeric variable by class. For Geomorphology classes, count of samples (and frequency, in percentage), and p-value from Chi-squared test.

| | **Damaged** | **Undamaged** | **p-value** |
|---|---|---|---|
| | ***N=5704*** | ***N=5800*** | |
| Tropical cyclone recurrence | 8.00 [8.00;8.00] | 8.00 [7.00;9.00] | 0.730 |
| Rainfall (dry season) (mm) | 135 [130;149] | 168 [85.3;205] | <0.001 |
| Rainfall (rainy season) (mm) | 1045 [832;1050] | 1105 [699;1296] | <0.001 |
| Wind speed (km/h) | 84.3 [65.7;89.4] | 56.0 [53.0;62.4] | 0.000 |
| Mean annual rainfall (mm) | 77.0 [70.8;128] | 118 [89.6;140] | 0.000 |
| Trend annual rainfall (mm/yr.) | 0.16 [0.15;0.16] | 0.14 [0.09;0.53] | <0.001 |
| Mean annual max. temperature (ºC) | 30.9 [30.0;31.6] | 31.8 [30.9;32.0] | 0.000 |
| Trend annual max. temperature (ºC/yr.) | 0.03 [0.03;0.04] | 0.03 [0.03;0.04] | <0.001 |
| Trend drought index (∆scPDSI/yr.) | -0.03 [-0.04;-0.03] | -0.04 [-0.06;-0.01] | 0.123 |
| Canopy height (m) | 10.2 [5.09;17.0] | 10.2 [5.09;17.0] | 0.012 |
| Green vegetation fractional cover | 0.81 [0.74;0.85] | 0.78 [0.70;0.84] | <0.001 |
| Soil fractional cover | 0.01 [0.00;0.03] | 0.02 [0.01;0.05] | <0.001 |
| Water fractional cover | 0.10 [0.08;0.12] | 0.09 [0.08;0.11] | <0.001 |
| Geomorphology class: | | | <0.001 |
| Delta | 775 (13.6%) | 2183 (37.6%) | |
| Estuary | 630 (11.0%) | 507 (8.74%) | |
| Lagoon | 4254 (74.6%) | 2860 (49.3%) | |
| OpenCoast | 45 (0.79%) | 250 (4.31%) | |
| Distance to shoreline (m) | 400 [141;762] | 791 [200;2983] | <0.001 |
| Distance to cropland (m) | 1709 [854;3177] | 3606 [1304;7362] | <0.001 |
| Distance to settlement (m) | 47997 [22489;54791] | 32628 [18883;44521] | <0.001 |
| Distance to road (m) | 6668 [2550;11413] | 5903 [2766;9962] | 0.036 |

**Appendix S14.** Vulnerability model data distribution from the Western Caribbean sub-region. Median [first; and third quartiles], and p-value from Kruskal-Wallis median test for each numeric variable by class. For Geomorphology classes, count of samples (and frequency, in percentage), and p-value from Chi-squared test.

| | **Damaged** *N=443* | **Undamaged** *N=450* | **p-value** |
|---|---|---|---|
| Tropical cyclone recurrence | 8.00 [8.00;9.00] | 8.00 [8.00;10.0] | <0.001 |
| Rainfall (dry season) (mm) | 419 [258;460] | 304 [227;439] | <0.001 |
| Rainfall (rainy season) (mm) | 1557 [1401;1934] | 1425 [1147;1694] | <0.001 |
| Wind speed (km/h) | 87.8 [53.8;95.5] | 55.7 [53.3;64.6] | <0.001 |
| Mean annual rainfall (mm) | 170 [147;170] | 147 [114;167] | <0.001 |
| Trend annual rainfall (mm/yr.) | 0.00 [-0.14;0.16] | 0.00 [-0.11;0.13] | 0.042 |
| Mean annual max. temperature (ºC) | 30.0 [29.5;30.0] | 30.0 [29.5;30.5] | <0.001 |
| Trend annual max. temperature (ºC/yr.) | 0.03 [0.02;0.03] | 0.03 [0.03;0.03] | <0.001 |
| Trend drought index (∆scPDSI/yr.) | -0.04 [-0.05;-0.02] | -0.04 [-0.05;-0.02] | 0.526 |
| Canopy height (m) | 8.48 [5.09;13.6] | 5.09 [3.39;8.48] | <0.001 |
| Green vegetation fractional cover | 0.82 [0.73;0.87] | 0.69 [0.58;0.77] | <0.001 |
| Soil fractional cover | 0.01 [0.00;0.02] | 0.04 [0.01;0.07] | <0.001 |
| Water fractional cover | 0.08 [0.06;0.12] | 0.12 [0.11;0.14] | <0.001 |
| Geomorphology class: | | | <0.001 |
| Delta | 4 (0.90%) | 0 (0.00%) | |
| Estuary | 23 (5.19%) | 27 (6.00%) | |
| Lagoon | 263 (59.4%) | 378 (84.0%) | |
| OpenCoast | 153 (34.5%) | 45 (10.0%) | |
| Distance to shoreline (m) | 200 [100;424] | 640 [200;1670] | <0.001 |
| Distance to cropland (m) | 1897 [600;18140] | 2892 [1211;6288] | 0.093 |
| Distance to settlement (m) | 16388 [14424;23017] | 30593 [18647;59107] | <0.001 |
| Distance to road (m) | 6555 [3135;18748] | 4545 [1905;7914] | <0.001 |

**Appendix S15.** The best number of predictors at each split in tree classification (mtry), and related overall accuracy (%) of each resilience random forest model.

| Resilience model | mtry | accuracy (%) |
|---|---|---|
| **Regional** | 15 | 87.27 |
| **Bahamian** | 9 | 83.39 |
| **Floridian** | 9 | 91.15 |
| **Greater Antilles** | 15 | 83.73 |
| **Southern Gulf of Mexico** | 15 | 80.44 |
| **Western Caribbean** | 21 | 74.12 |

**Appendix S16.** Resilience model data distribution from the entire North Atlantic Basin region. Median [first; and third quartiles], and p-value from Kruskal-Wallis median test for each numeric variable by class. For Geomorphology classes, count of samples (and frequency, in percentage), and p-value from Chi-squared test.

| | **Lost** *N=59109* | **Recovered** *N=59322* | **p-value** |
|---|---|---|---|
| Tropical cyclone recurrence | 8.00 [7.00;9.00] | 8.00 [7.00;9.00] | <0.001 |
| Rainfall (dry season) (mm) | 171 [135;216] | 143 [124;196] | 0.000 |
| Rainfall (rainy season) (mm) | 1052 [985;1194] | 1043 [898;1205] | <0.001 |
| Wind speed (km/h) | 104 [90.9;115] | 108 [93.6;123] | 0.000 |
| Mean annual rainfall (mm) | 105 [100;112] | 105 [100;116] | <0.001 |
| Trend annual rainfall (mm/yr.) | 0.21 [0.12;0.33] | 0.32 [0.16;0.33] | 0.000 |
| Mean annual max. temperature (ºC) | 29.8 [28.6;30.4] | 29.2 [28.6;30.0] | <0.001 |
| Trend annual max. temperature (ºC/yr.) | 0.02 [0.01;0.02] | 0.01 [0.01;0.02] | 0.000 |
| Trend drought index (∆scPDSI/yr.) | -0.02 [-0.04;0.04] | 0.03 [-0.02;0.04] | 0.000 |
| Canopy height (m) | 8.48 [5.09;13.6] | 13.6 [8.48;18.7] | 0.000 |
| Green vegetation fractional cover | 0.81 [0.70;0.88] | 0.86 [0.79;0.90] | 0.000 |
| Soil fractional cover | 0.01 [0.00;0.03] | 0.01 [0.00;0.02] | 0.000 |
| Water fractional cover | 0.07 [0.04;0.10] | 0.07 [0.04;0.09] | <0.001 |
| Geomorphology class: | | | <0.001 |
| Delta | 1058 (1.79%) | 737 (1.24%) | |
| Estuary | 704 (1.19%) | 1557 (2.62%) | |
| Lagoon | 41392 (70.0%) | 44453 (74.9%) | |
| OpenCoast | 15955 (27.0%) | 12575 (21.2%) | |
| Soil organic carbon stock (Mg C/ ha) | 509 [404;554] | 522 [455;564] | <0.001 |
| Distance to shoreline (m) | 671 [200;1836] | 707 [224;1628] | 0.977 |
| Distance to cropland (m) | 12146 [2561;41109] | 12902 [1942;54403] | <0.001 |
| Distance to settlement (m) | 28231 [15940;40806] | 28037 [14031;47324] | <0.001 |
| Distance to road (m) | 9552 [4662;17806] | 8792 [3920;23456] | 0.084 |

**Appendix S17.** Resilience model data distribution from the Bahamian sub-region. Median [first; and third quartiles], and p-value from Kruskal-Wallis median test for each numeric variable by class. For Geomorphology classes, count of samples (and frequency, in percentage), and p-value from Chi-squared test.

| | **Lost** *N=2794* | **Recovered** *N=2725* | **p-value** |
|---|---|---|---|
| Tropical cyclone recurrence | 9.00 [8.00;9.00] | 9.00 [8.00;9.00] | <0.001 |
| Rainfall (dry season) (mm) | 301 [294;370] | 286 [219;358] | <0.001 |
| Rainfall (rainy season) (mm) | 1335 [1124;1420] | 1124 [1031;1335] | <0.001 |
| Wind speed (km/h) | 110 [90.9;115] | 115 [99.4;127] | <0.001 |
| Mean annual rainfall (mm) | 105 [104;105] | 105 [102;105] | 0.454 |
| Trend annual rainfall (mm/yr.) | -0.06 [-0.10;0.01] | -0.05 [-0.10;0.04] | <0.001 |
| Mean annual max. temperature (ºC) | 28.5 [28.5;28.5] | 28.5 [28.5;28.8] | 0.127 |
| Trend annual max. temperature (ºC/yr.) | 0.02 [0.02;0.02] | 0.02 [0.02;0.02] | <0.001 |
| Trend drought index (∆scPDSI/yr.) | -0.07 [-0.07;-0.07] | -0.07 [-0.07;-0.06] | <0.001 |
| Canopy height (m) | 1.70 [0.85;3.39] | 1.70 [0.85;5.09] | <0.001 |
| Green vegetation fractional cover | 0.53 [0.42;0.65] | 0.58 [0.47;0.69] | <0.001 |
| Soil fractional cover | 0.02 [0.01;0.06] | 0.02 [0.01;0.06] | <0.001 |
| Water fractional cover | 0.15 [0.08;0.23] | 0.14 [0.08;0.22] | 0.063 |
| Geomorphology class: | | | <0.001 |
| Lagoon | 64 (2.29%) | 150 (5.50%) | |
| OpenCoast | 2730 (97.7%) | 2575 (94.5%) | |
| Distance to shoreline (m) | 224 [100;1393] | 224 [100;1281] | 0.336 |
| Distance to cropland (m) | 12361 [8160;35878] | 18913 [10689;48194] | <0.001 |
| Distance to settlement (m) | 22310 [13653;27259] | 24512 [12078;30973] | <0.001 |
| Distance to road (m) | 8222 [5200;11679] | 9546 [5590;14584] | <0.001 |
| Soil organic carbon stock (Mg C/ ha) | 408 [368;516] | 452 [382;533] | <0.001 |

**Appendix S18.** Resilience model data distribution from the Floridian sub-region. Median [first; and third quartiles], and p-value from Kruskal-Wallis median test for each numeric variable by class. For Geomorphology classes, count of samples (and frequency, in percentage), and p-value from Chi-squared test.

| | **Lost** | **Recovered** | **p-value** |
|---|---|---|---|
| | ***N=18112*** | ***N=18017*** | |
| Tropical cyclone recurrence | 9.00 [7.00;9.00] | 9.00 [8.00;9.00] | <0.001 |
| Rainfall (dry season) (mm) | 144 [128;162] | 133 [123;147] | <0.001 |
| Rainfall (rainy season) (mm) | 1052 [1007;1225] | 1043 [966;1225] | 0.101 |
| Wind speed (km/h) | 106 [93.1;108] | 110 [105;128] | 0.000 |
| Mean annual rainfall (mm) | 112 [100;112] | 102 [100;112] | <0.001 |
| Trend annual rainfall (mm/yr.) | 0.51 [0.32;0.55] | 0.33 [0.32;0.51] | <0.001 |
| Mean annual max. temperature (ºC) | 28.6 [28.6;29.2] | 28.8 [28.6;29.2] | <0.001 |
| Trend annual max. temperature (ºC/yr.) | 0.01 [0.01;0.01] | 0.01 [0.01;0.01] | <0.001 |
| Trend drought index (∆scPDSI/yr.) | 0.04 [0.04;0.06] | 0.04 [0.04;0.04] | <0.001 |
| Canopy height (m) | 11.9 [8.48;15.3] | 15.3 [11.9;20.4] | 0.000 |
| Green vegetation fractional cover | 0.84 [0.78;0.89] | 0.86 [0.81;0.90] | <0.001 |
| Soil fractional cover | 0.01 [0.00;0.02] | 0.00 [0.00;0.01] | 0.000 |
| Water fractional cover | 0.06 [0.04;0.08] | 0.07 [0.04;0.08] | <0.001 |
| Geomorphology class: | | | <0.001 |
| Estuary | 92 (0.51%) | 350 (1.94%) | |
| Lagoon | 16431 (90.7%) | 16415 (91.1%) | |
| OpenCoast | 1589 (8.77%) | 1252 (6.95%) | |
| Distance to shoreline (m) | 1204 [400;2774] | 1030 [400;2220] | <0.001 |
| Distance to cropland (m) | 54127 [38492;62150] | 53668 [23715;61621] | <0.001 |
| Distance to settlement (m) | 33912 [13415;45254] | 42177 [11555;55439] | <0.001 |
| Distance to road (m) | 13788 [5914;28800] | 16805 [5457;31621] | <0.001 |
| Soil organic carbon stock (Mg C/ ha) | 549 [527;567] | 552 [525;581] | <0.001 |

**Appendix S19.** Resilience model data distribution from the Greater Antilles sub-region. Median [first; and third quartiles], and p-value from Kruskal-Wallis median test for each numeric variable by class. For Geomorphology classes, count of samples (and frequency, in percentage), and p-value from Chi-squared test.

| | **Lost** | **Recovered** | **p-value** |
|---|---|---|---|
| | ***N=21095*** | ***N=21033*** | |
| Tropical cyclone recurrence | 8.00 [7.00;9.00] | 8.00 [6.00;9.00] | <0.001 |
| Rainfall (dry season) (mm) | 203 [167;216] | 191 [137;219] | <0.001 |
| Rainfall (rainy season) (mm) | 1024 [913;1133] | 1037 [870;1167] | <0.001 |
| Wind speed (km/h) | 107 [95.5;118] | 104 [91.3;116] | <0.001 |
| Mean annual rainfall (mm) | 105 [101;121] | 109 [105;123] | 0.000 |
| Trend annual rainfall (mm/yr.) | 0.20 [0.13;0.26] | 0.21 [0.16;0.28] | <0.001 |
| Mean annual max. temperature (ºC) | 30.1 [29.8;30.5] | 30.0 [29.8;30.4] | <0.001 |
| Trend annual max. temperature (ºC/yr.) | 0.02 [0.01;0.02] | 0.02 [0.01;0.02] | <0.001 |
| Trend drought index (∆scPDSI/yr.) | -0.02 [-0.04;-0.02] | -0.02 [-0.04;0.02] | <0.001 |
| Canopy height (m) | 8.48 [5.09;13.6] | 13.6 [8.48;17.0] | 0.000 |
| Green vegetation fractional cover | 0.84 [0.74;0.91] | 0.88 [0.82;0.93] | <0.001 |
| Soil fractional cover | 0.01 [0.00;0.02] | 0.01 [0.00;0.02] | <0.001 |
| Water fractional cover | 0.05 [0.02;0.08] | 0.04 [0.02;0.08] | <0.001 |
| Geomorphology class: | | | <0.001 |
| Delta | 25 (0.12%) | 6 (0.03%) | |
| Estuary | 28 (0.13%) | 56 (0.27%) | |
| Lagoon | 16371 (77.6%) | 13083 (62.2%) | |
| OpenCoast | 4671 (22.1%) | 7888 (37.5%) | |
| Distance to shoreline (m) | 700 [224;1655] | 510 [200;1140] | <0.001 |
| Distance to cropland (m) | 5201 [1500;14849] | 1844 [1000;4950] | 0.000 |
| Distance to settlement (m) | 26443 [16206;35740] | 22699 [14825;29659] | <0.001 |
| Distance to road (m) | 10519 [4245;18776] | 5536 [3008;11328] | 0.000 |
| Soil organic carbon stock (Mg C/ ha) | 505 [463;551] | 491 [441;529] | <0.001 |

**Appendix S20.** Resilience model data distribution from the Southern Gulf of Mexico sub-region. Median [first; and third quartiles], and p-value from Kruskal-Wallis median test for each numeric variable by class. For Geomorphology classes, count of samples (and frequency, in percentage), and p-value from Chi-squared test.

| | **Lost** | **Recovered** | **p-value** |
|---|---|---|---|
| | ***N=5387*** | ***N=5352*** | |
| Tropical cyclone recurrence | 8.00 [8.00;8.00] | 8.00 [8.00;8.00] | 0.088 |
| Rainfall (dry season) (mm) | 135 [130;149] | 135 [125;149] | <0.001 |
| Rainfall (rainy season) (mm) | 1045 [857;1050] | 1036 [822;1050] | 0.009 |
| Wind speed (km/h) | 85.8 [66.8;89.4] | 70.6 [53.9;89.4] | <0.001 |
| Mean annual rainfall (mm) | 77.0 [70.8;77.0] | 70.8 [70.8;128] | 0.023 |
| Trend annual rainfall (mm/yr.) | 0.15 [0.15;0.16] | 0.16 [0.14;0.16] | <0.001 |
| Mean annual max. temperature (ºC) | 30.9 [30.0;31.6] | 30.3 [30.0;31.6] | <0.001 |
| Trend annual max. temperature (ºC/yr.) | 0.03 [0.03;0.04] | 0.03 [0.03;0.04] | <0.001 |
| Trend drought index (∆scPDSI/yr.) | -0.04 [-0.04;-0.03] | -0.03 [-0.05;-0.03] | <0.001 |
| Canopy height (m) | 10.2 [5.09;17.0] | 11.9 [6.79;17.0] | <0.001 |
| Green vegetation fractional cover | 0.80 [0.74;0.85] | 0.80 [0.72;0.85] | <0.001 |
| Soil fractional cover | 0.01 [0.00;0.04] | 0.01 [0.00;0.04] | 0.076 |
| Water fractional cover | 0.10 [0.08;0.12] | 0.10 [0.09;0.13] | <0.001 |
| Geomorphology class: | | | <0.001 |
| Delta | 756 (14.0%) | 692 (12.9%) | |
| Estuary | 406 (7.54%) | 873 (16.3%) | |
| Lagoon | 4185 (77.7%) | 3751 (70.1%) | |
| OpenCoast | 40 (0.74%) | 36 (0.67%) | |
| Distance to shoreline (m) | 412 [141;781] | 400 [141;800] | 0.055 |
| Distance to cropland (m) | 1803 [985;3158] | 1513 [671;3162] | <0.001 |
| Distance to settlement (m) | 49786 [26998;55109] | 44398 [19496;54101] | <0.001 |
| Distance to road (m) | 6325 [2550;10026] | 7750 [2467;12142] | <0.001 |
| Soil organic carbon stock (Mg C/ ha) | 284 [234;337] | 268 [200;337] | <0.001 |

**Appendix S21.** Resilience model data distribution from the Western Caribbean sub-region. Median [first; and third quartiles], and p-value from Kruskal-Wallis median test for each numeric variable by class. For Geomorphology classes, count of samples (and frequency, in percentage), and p-value from Chi-squared test.

| | **Lost** | **Recovered** | **p-value** |
|---|---|---|---|
| | ***N=235*** | ***N=221*** | |
| Tropical cyclone recurrence | 8.00 [8.00;9.00] | 8.00 [8.00;8.00] | 0.710 |
| Rainfall (dry season) (mm) | 381 [248;460] | 460 [434;474] | <0.001 |
| Rainfall (rainy season) (mm) | 1557 [1447;1984] | 1557 [1489;1686] | 0.002 |
| Wind speed (km/h) | 87.8 [50.1;99.8] | 95.5 [53.8;95.5] | 0.587 |
| Mean annual rainfall (mm) | 147 [147;170] | 170 [170;170] | <0.001 |
| Trend annual rainfall (mm/yr.) | 0.00 [-0.14;0.16] | 0.16 [0.00;0.16] | <0.001 |
| Mean annual max. temperature (ºC) | 30.0 [29.5;30.0] | 29.5 [29.5;30.0] | <0.001 |
| Trend annual max. temperature (ºC/yr.) | 0.03 [0.02;0.03] | 0.02 [0.02;0.03] | <0.001 |
| Trend drought index (∆scPDSI/yr.) | -0.05 [-0.05;-0.02] | -0.02 [-0.04;-0.02] | <0.001 |
| Canopy height (m) | 8.48 [5.09;13.6] | 8.48 [5.09;13.6] | 0.114 |
| Green vegetation fractional cover | 0.83 [0.75;0.87] | 0.84 [0.77;0.87] | 0.086 |
| Soil fractional cover | 0.01 [0.00;0.02] | 0.01 [0.00;0.02] | 0.281 |
| Water fractional cover | 0.09 [0.07;0.12] | 0.07 [0.06;0.10] | <0.001 |
| Geomorphology class: | | | <0.001 |
| Delta | 1 (0.43%) | 8 (3.62%) | |
| Estuary | 5 (2.13%) | 24 (10.9%) | |
| Lagoon | 133 (56.6%) | 164 (74.2%) | |
| OpenCoast | 96 (40.9%) | 25 (11.3%) | |
| Distance to shoreline (m) | 141 [100;418] | 141 [100;300] | 0.095 |
| Distance to cropland (m) | 2751 [632;18429] | 900 [361;4000] | <0.001 |
| Distance to settlement (m) | 16629 [14603;22962] | 16571 [14144;22402] | 0.711 |
| Distance to road (m) | 6986 [4387;21004] | 6522 [3311;7739] | 0.009 |
| Soil organic carbon stock (Mg C/ ha) | 402 [346;457] | 448 [370;471] | 0.001 |
